# The cellular basis of mitochondrial stress signaling in the brain

**DOI:** 10.1101/2025.09.11.675336

**Authors:** Emma L. Hamer, Vincent Gardeux, Radiana Ferrero, Bart Deplancke, Joseph M. Bateman

**Affiliations:** Maurice Wohl Clinical Neuroscience Institute; King’s College London, London, SE5 9RX, UK; Laboratory of Systems Biology and Genetics; Institute of Bioengineering, School of Life Sciences, Ecole Polytechnique Fédérale de Lausanne (EPFL), CH-1015 Lausanne, Switzerland; Swiss Institute of Bioinformatics; CH-1015 Lausanne, Switzerland

## Abstract

Neuronal vulnerability to stress is highly cell-type specific, but the underlying mechanisms are poorly understood. Here we show that the ability to activate endoplasmic reticulum (ER) stress signaling is encoded by the relative mitochondrial metabolic activity of neurons. Genetically inducing mitochondrial dysfunction in the *Drosophila* brain caused the emergence of a novel cluster of neurons, detected using single nuclear RNA-sequencing, that activated the ER unfolded protein response (UPR). UPR activation occurred only in neurons with high mitochondrial activity. Unexpectedly, mitochondria-ER contacts were also abundant in the neurons that activated the UPR but virtually undetectable in neurons with low mitochondrial activity. In the human brain, strikingly, excitatory neurogranin neurons had the highest mitochondrial activity and triggered the UPR. The selective activation of the UPR only in neurons with high mitochondrial activity is therefore conserved. This study reveals the remarkable dependence on the inherent oxidative metabolic activity of neurons to activate the UPR.

## Main

Adaptation to changing environmental conditions to maintain homeostasis is a key feature of eukaryotic cells. The ability to modify the cellular environment under stressful conditions is an adaptive response that helps to maintain cellular and organismal viability, avoiding cell death^1^. The brain is composed of myriad neuron types each with specific morphologies, functions and network connections^2^. The distinct cellular and molecular properties of each type of neuron are essential for healthy brain function^3^. Cellular stress and disease pathology do not affect all neuron types equally, but the mechanisms underlying the cell-type specificity of the response to cellular stress in the brain are unknown.

Mitochondria act as signaling organelles. Perturbed mitochondrial function, due to, for example, nutrient deprivation, senescence, or disease, trigger mitochondrial stress signaling pathways^4, 5^. The mechanism of mitochondrial stress signaling depends on the cellular context, but the most frequently activated signaling pathway is the endoplasmic reticulum (ER) unfolded protein response (UPR)^4, 6, 7^. The UPR is triggered by the loss of ER proteostasis and consists of a coordinated transcriptional response mediated by the ER transmembrane kinase PERK, inositol-requiring enzyme 1 (IRE1) and activating transcription factor 6 (ATF6)^8, 9^. The UPR regulates the expression of genes that coordinate an adaptive response to reduce ER stress, or alternatively trigger apoptosis.

Compromised mitochondrial function causes rare primary mitochondrial diseases and is a hallmark of common neurological diseases including Parkinson’s disease (PD) and Alzheimer’s disease (AD), autism spectrum disorders and schizophrenia^10, 11^. Stress signaling pathways, in particular the UPR, are activated in animal models of primary mitochondrial diseases, PD, AD and frontotemporal dementia^6, 12–15^. However, we have very limited understanding of how the wide range of cell types in the brain respond to mitochondrial dysfunction, or whether the UPR is activated constitutively, or in specific cell types in the brain. We used single nuclear RNA-seq (snRNA-seq) of a *Drosophila* model of neuronal mitochondrial dysfunction and snRNA-seq data from the human brain to identify the cell types that activate the UPR. Our study reveals a general principle, that UPR activation is determined by neuronal mitochondrial activity.

## Results

### snRNA-seq reveals mitochondrial dysfunction causes transcriptional reprogramming of neurons

We used knockdown of the *Drosophila* NDUFS1 homolog ND-75 (ND-75 KD), a core catalytic subunit of complex I of the respiratory chain^16, 17^, to model the effects of mitochondrial dysfunction in neurons and dissect the mitochondrial stress signaling mechanisms in the brain. We performed snRNA-seq of heads from flies with pan-neuronal ND-75 KD and controls and obtained transcriptomes from 11,437 control cells and 11,742 ND-75 KD cells (Fig. 1A and Methods). After integration, we found 90 distinct clusters represented by Uniform Manifold Approximation and Projection (UMAP) (Fig. 1B; Supplementary Fig. S1A). Using previous high depth single cell RNA-seq and snRNA-seq data sets^18, 19^, we annotated these clusters to identifiable classes of neurons and non-neuronal cells. Using the integrated data set we annotated 15,165 cells representing 50 different cell types leaving 8,014 unannotated cells, a similar proportion to previous studies of *Drosophila* brain and head tissue^18, 19^. These annotated cell types represented the major classes of neurons and glia, as well as non-CNS cells including photoreceptors, head muscle cells and hemocytes (Fig. 1B, Supplementary Fig. S1A). Most of the clusters that were detected across the two datasets were reproduced with high transcriptomic similarity, which is highlighted in the integrated UMAP of control and ND-75 KD cells (Fig. 1C), even without Harmony integration (Supplementary Fig. S1B).

**Fig. 1.**
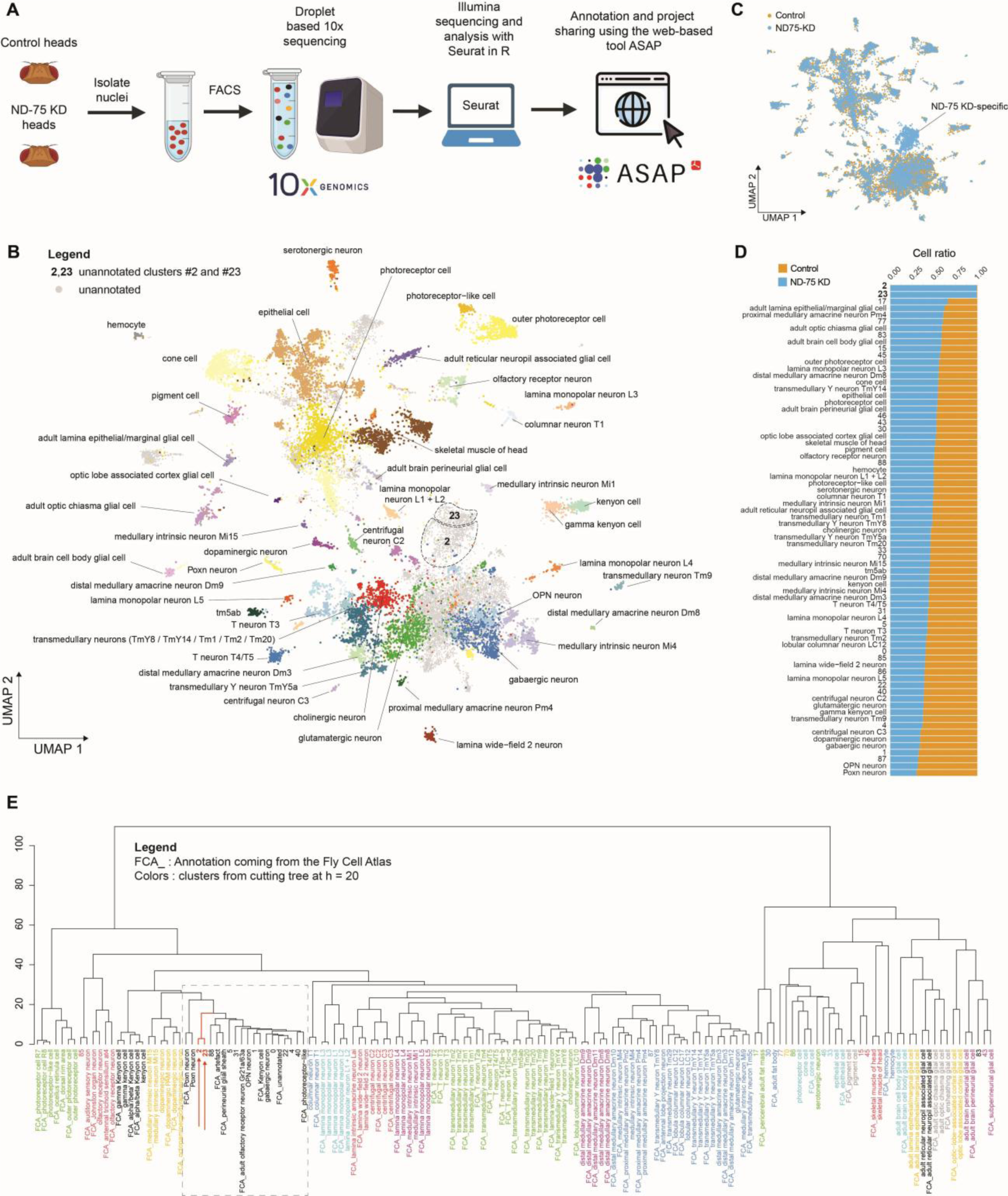
ND-75 KD transcriptionally reprogrammes a subset of neurons in the Drosophila brain. (A) Schematic of the snRNA-seq pipeline. (B) UMAP showing annotated colour coded cell clusters in the integrated snRNA-seq dataset. (C) UMAP showing control (orange) and ND-75 KD (blue) cell identities with Harmony integration. Novel clusters 2 and 23 consist almost exclusively of ND-75 KD cells. (D) Chart showing the contribution of control (orange) and ND-75 KD (blue) cells to each cluster (numbered clusters are unannotated). Clusters 2 and 23 consist almost entirely of ND-75 KD cells. (E) Hierarchical clustering (Euclidean distance, ward.D2 clustering method) computed on the Harmony-integrated dataset between the control and ND-75 KD integrated dataset and the Fly Cell Atlas dataset. Dendrogram of the annotated clusters shows clusters 2 and 23 (arrows) are closely related to Poxn neurons, OPNs and several other types of neurons.

Although the clusters were largely equivalent between control and ND-75 KD conditions, two clusters, 2 and 23, consisting of 817 and 333 cells respectively, emerged almost exclusively in ND-75 KD flies (Fig. 1C, D; Supplementary Fig. S1B). Clusters 2 and 23 therefore represent transcriptionally novel cells that are reprogrammed due to ND-75 KD. Comparison of the marker genes in clusters 2 and 23 with head tissue marker genes from FlyCellAtlas^19^ showed that very few cell-type marker genes were present in the novel clusters (Supplementary Tables S1, S2), consistent with the UMAP (Fig. 1C), indicating that the transcriptional reprogramming of these cells masks their identity. The novel clusters could potentially derive from a single cell type that is highly sensitive to mitochondrial dysfunction, or from multiple cell types that converge to a common novel transcriptional identity. Hierarchical clustering of the integrated data together with the Fly Cell Atlas dataset^19^, showed that clusters 2 and 23 were closely associated with multiple neuron types including olfactory receptor Gr21a/63a neurons, Kenyon cells, GABAergic neurons, olfactory projection neurons (OPNs) and Poxn neurons (Fig. 1E). Consistent with this, in the integrated dataset OPNs and Poxn neuron clusters had the lowest number of ND-75 KD cells (Fig. 1D). These data may indicate that novel clusters 2 and 23 consist of multiple types of neurons, including OPNs and Poxn neurons, that have been transcriptionally reprogrammed by ND-75 KD.

### Mitochondrial dysfunction activates the UPR in a subset of central brain neurons

Mitochondrial dysfunction activates mitochondrial stress signaling pathways that employ key transcription factors to impose large-scale changes in gene expression^6^. To identify transcriptional regulators involved in the reprogramming of neurons caused by ND-75 KD, we predicted gene regulatory networks based on co-expression and motif enrichment using SCENIC^20^. Then, we used this regulon space to generate a “regulon-based UMAP”, resulting in a different UMAP projection based on transcription factor regulon activity, which showed again a clear separation of clusters 2 and 23 (Fig. 2A). Differential regulon activity analysis showed that several transcription factors had significantly increased activity in both clusters 2 and 23 including UPR components Atf6 and crc, the *Drosophila* homolog of ATF4 (Fig 2B-D)^9^. ATF4 is a highly conserved basic leucine zipper superfamily transcription factor that is activated by the UPR (Fig. 2D)^7^. After photoreceptors, which normally express *crc* during development^21, 22^, *crc* was most highly expressed in clusters 2 and 23 (Fig. 2E). Binarised regulon-based UMAP representation and direct comparison showed that the crc regulon had significantly higher enrichment in cluster 23 than cluster 2 (Fig. 2F, G). Thus, the enrichment of UPR-related transcription factor activity in clusters 2 and 23 suggests that the UPR contributes to mitochondrial stress signaling and transcriptional reprogramming of *Drosophila* neurons.

**Fig. 2.**
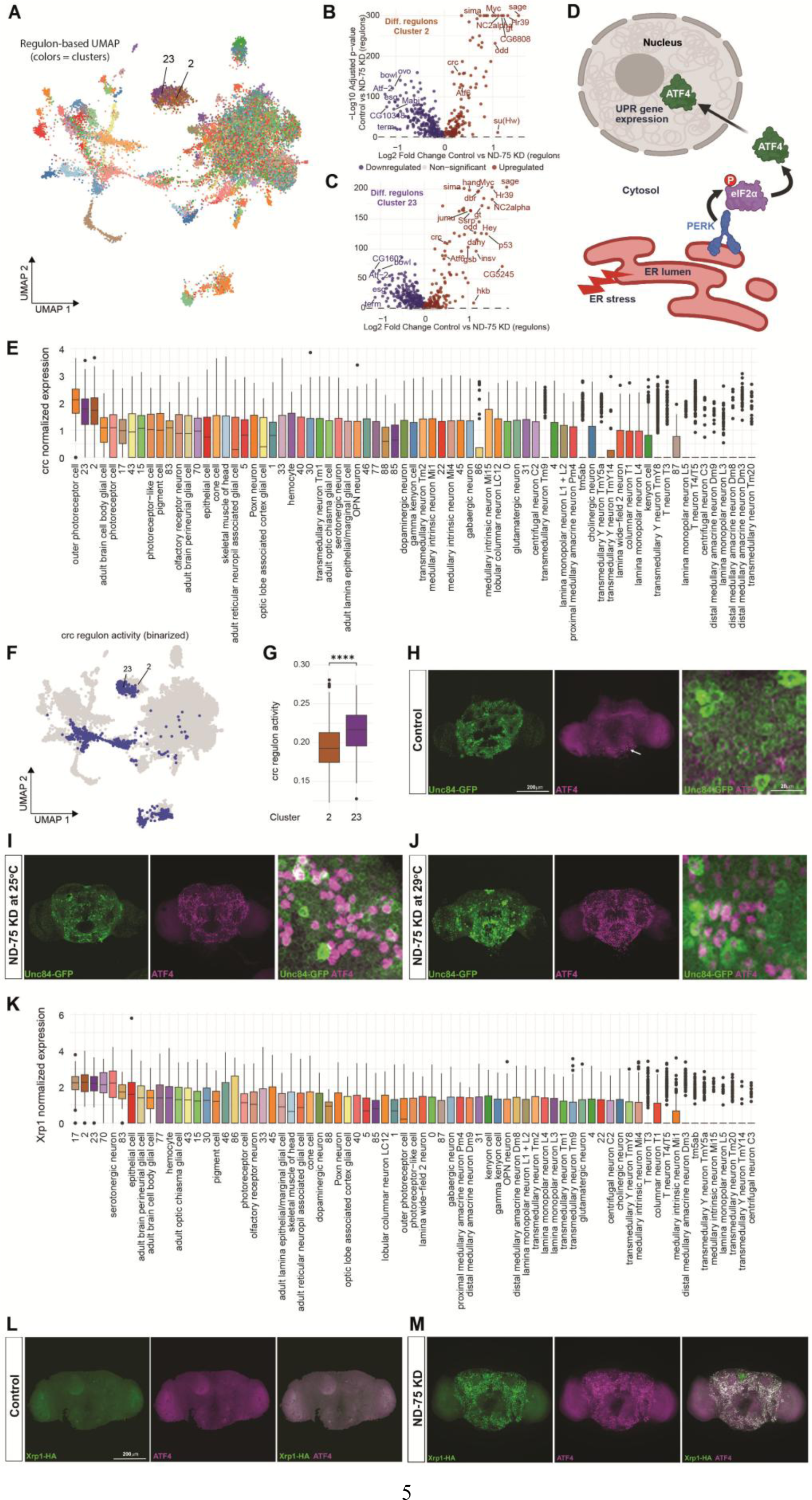
ATF4 and Xrp1 are enriched in the novel clusters and activated by ND-75 KD in a specific subset of neurons. (A) Regulon-based UMAP of the control and ND-75 KD integrated snRNA-seq datasets. Clusters are colour coded using the same colouring pattern defined in Supplementary Figure S1A. Novel clusters 2 and 23 are labelled. (B, C) Significantly increased transcription factor regulons in clusters 2 (B) and 23 (C) using Seurat’s Wilcoxon’s test on the regulon enrichment matrix. (D) Schematic of ER stress activating the UPR and ATF4. (E) *crc* is highly expressed in clusters 2 and 23 shown in ranked clusters. (F) Regulon-based binarised UMAP of crc regulated genes showing enrichment in clusters 2 and 23. (G) crc regulon activity is significantly higher in cluster 23 than cluster 2 according to Wilcoxon rank-sum test. (H-J) ATF4 (magenta) is only expressed in a small number of cells in the suboesophageal ganglion (arrow) in *Tub-Gal80^ts^;nSyb-Gal4*-driven luciferase RNAi control brain (H), but is strongly activated throughout the central brain in a specific subset of neurons with pan-neuronal (*Tub-Gal80^ts^;nSyb-Gal4*) ND-75 KD at 25°C (I) and 29°C (J, late pupal brain). Unc84-GFP (green) expression is shown as a reporter of *nSyb-Gal4* activity. Whole brain images are maximum intensity projections. Right hand close-up panels are single z-plane images. (K) *Xrp1* is highly expressed in clusters 2 and 23 shown in ranked clusters. (L) Pan-neuronal (*Tub-Gal80^ts^;nSyb-Gal4*-driven) luciferase RNAi does not activate Xrp1-HA (green) or ATF4 (magenta) in the brain. (M) Pan-neuronal (*Tub-Gal80^ts^;nSyb-Gal4*) ND-75 KD at 25°C strongly activates Xrp1-HA (green) and ATF4 (magenta) in the same neurons. All brain images apart from (J) are from 1-2 day old flies. ****p < 0.0001.

To further examine the cell-type specific activation of ATF4 in ND-75 KD neurons we performed immunostaining. In control adult brains ATF4 was expressed in on average only 29 cells (±2.7, n=5 brains) in the suboesophageal ganglion (Fig. 2H, arrow; Supplementary Fig. S2A, arrow). Pan-neuronal ND-75 KD using *nSyb-Gal4* caused pupal lethality but adults were viable using *Tub-Gal80^ts^;nSyb-Gal4* at 25°C^17^. ND-75 KD using *Tub-Gal80^ts^;nSyb-Gal4* at 25°C caused ATF4 expression in on average 1099 neurons (±122, n=5 adult brains) in the central brain (Fig. 2I, Supplementary Fig. S2B). Under conditions of maximum Gal4 activity, using *Tub-Gal80^ts^;nSyb-Gal4* to knockdown ND-75 at 29°C, which caused late pupal lethality, pan-neuronal ND-75 KD caused ATF4 activation in on average 1527 neurons (±93, n=5 late pupal brains) (Fig. 2J). Pan-neuronal mitochondrial dysfunction therefore caused ATF4 activation in around 1% of the 140,000 neurons in the *Drosophila* brain^23^. Neurons that failed to activate ATF4 were not a result of lower *nSyb-Gal4* activity since, using nuclear GFP as a marker of Gal4 activity, ND-75 KD caused ATF4 activation in neurons with both high and low Gal4 activity (Fig. 2I, J, right hand panels). The pattern of ATF4 activation was also independent of the *Gal4* driver, as a very similar pattern of ATF4 activation was seen with pan-neuronal ND-75 KD using *Appl-Gal4;Tub-Gal80^ts^* at 25°C (Supplementary Fig. S2C, D).

To test whether an independent method of inducing mitochondrial dysfunction activates ATF4 in a similar spatial pattern to ND-75 KD, we used overexpression of the inner mitochondrial membrane dynamin-related GTPase Opa1. Opa1 regulates mitochondrial dynamics, integrity and mitochondrial DNA maintenance and overexpression of Opa1 in *Drosophila* neurons causes mitochondrial fragmentation and mitochondrial dysfunction^24^. Pan-neuronal Opa1 overexpression using *Tub-Gal80^ts^;nSyb-Gal4* at 29°C caused late pupal lethality and a very similar pattern of ATF4 activation in neurons in the central brain to ND-75 KD (Supplementary Fig. S2E, F). Thus, pan-neuronal mitochondrial dysfunction activates ATF4 in a specific subset of neurons in the central brain.

We next looked for additional UPR components with high expression in clusters 2 and 23 to validate the findings with ATF4. Similar to ATF4, expression of the AT-hook bZip DNA binding protein Xrp1 is regulated by the UPR^25, 26^ and Xrp1 was very highly expressed in clusters 2 and 23 (Fig. 2K). By analysing the expression of an endogenously HA-tagged allele of Xrp1^26^, we found that Xrp1-HA was not expressed in control brains, but pan-neuronal ND-75 KD caused strong activation of Xrp1-HA expression in the same neurons that activated ATF4 (Fig. 2L, M). These data further demonstrate that mitochondrial dysfunction activates the UPR in a specific subset of neurons in the *Drosophila* central brain.

### Mitochondrial dysfunction activates ATF4 in central brain B, Imp-expressing neurons

We next investigated the basis for the ND-75 KD-dependent activation of ATF4 in a specific subset of neurons in the *Drosophila* central brain. To test whether the neuron specific activation of ATF4 was dependent on neurotransmitter type, we used cell-type-specific Gal4 drivers to perform ND-75 KD in cholinergic, glutamatergic and GABAergic neurons. ND-75 KD caused ATF4 activation in all three types of neurons (Supplementary Fig. S3A-F), demonstrating that neurotransmitter identity does not determine which neurons activate ATF4.

Since ATF4 activation was not associated with a specific type of neurotransmitter and cluster 2 and 23 were closely related to a range of different neuron types, we looked for alternative explanations for the activation of the UPR in a specific subset of neurons. *Drosophila* neurons can be categorised into three classes based on the transcription factors Prospero (Pros) and datilografo (dati; central brain A neurons), IGF-II mRNA-binding protein (Imp; central brain B neurons) and scarecrow (scro; optic lobe neurons)^18^. Cluster 23 was enriched for central brain B neurons (Fig. 3A), consistent with cluster 23 scoring very high for Imp expression but much lower for Pros and Dati expression (Supplementary Fig. S3G-I). Cluster 23 also had the highest level of ATF4 regulon activity (Fig. 2F, G). The intersection of these data suggested that ATF4 may be activated in central brain B neurons. To corroborate these snRNA-seq data, we analysed the expression of Pros and Imp in ND-75 KD brains. Pan-neuronal ND-75 KD activated ATF4 expression in Imp-expressing neurons (Fig. 3B), but not in the majority of Pros-expressing neurons (Fig. 3C). Moreover, activation of ATF4 by inducing mitochondrial dysfunction using overexpression of Opa1 resulted in the same pattern of ATF4 expression in Imp-expressing neurons but not most Pros-expressing neurons (Fig. 3D, E). Therefore, mitochondrial dysfunction activates the UPR in Imp-expressing central brain B neurons.

**Fig. 3.**
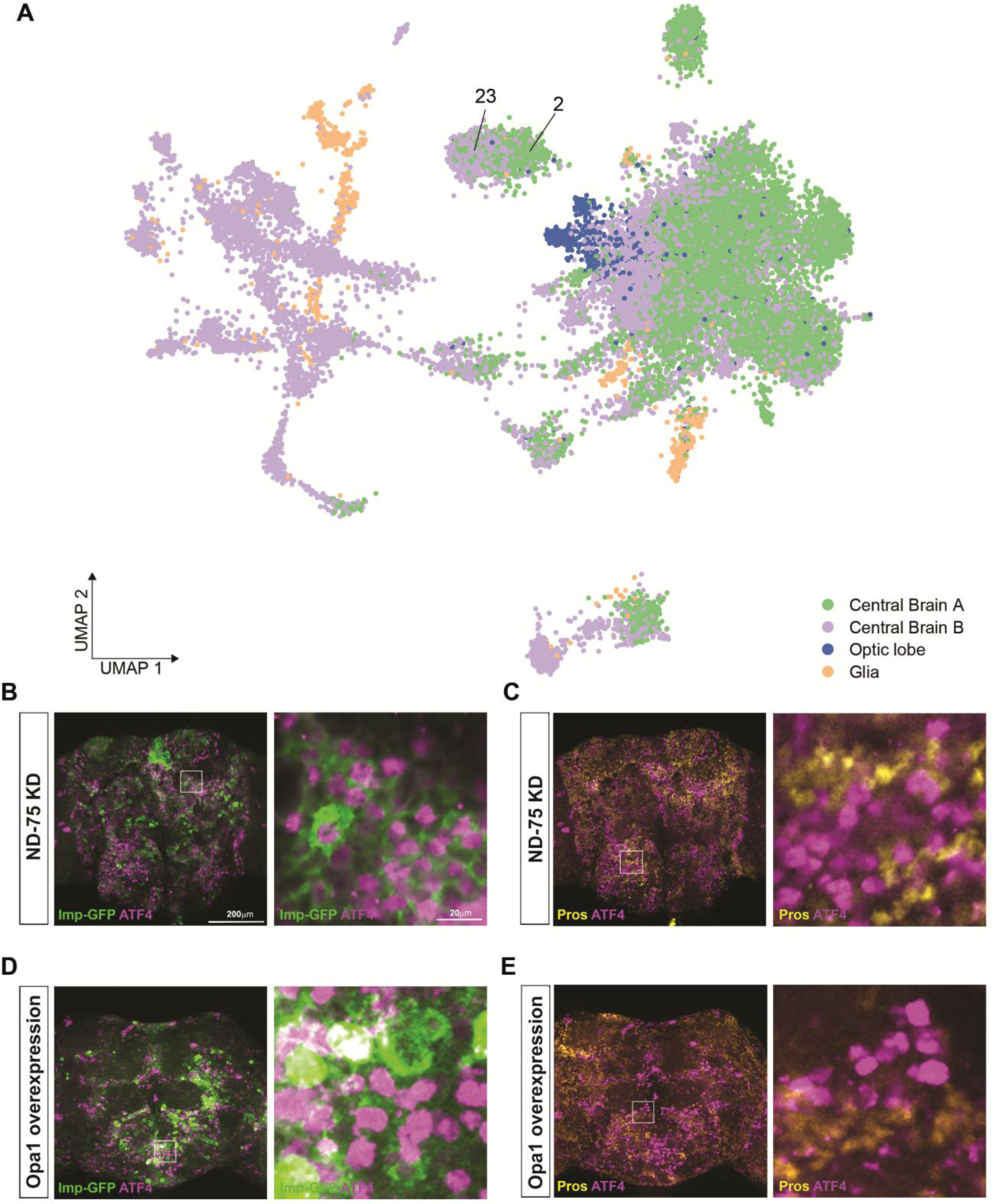
Mitochondrial dysfunction activates ATF4 in central brain B neurons. (A) Binarised regulon-UMAP showing cells coloured by cell type categorization, using Pros/dati (central brain A neurons), scro (optic lobe neurons), Repo (glia) regulon binarized activity and the remaining cells as central brain B neurons (see Methods). Novel cluster 23 is enriched for central brain B neurons. (B, C) Pan-neuronal (*Tub-Gal80ts;nSyb-Gal4*) ND-75 KD at 25°C activates ATF4 (magenta) in neurons that express Imp-GFP (localised in the cytosol, green, B) but not in neurons that express Pros (nuclear, yellow, C). Images are from 1-2 day old adult flies. (D, E) Pan-neuronal (*Tub-Gal80ts;nSyb-Gal4*) Opa1 overexpression at 25°C activates ATF4 (magenta) in neurons that express Imp-GFP (localised in the cytosol, green, D) but not in neurons that express Pros (nuclear, yellow, E) in the late pupal brain. White boxes indicate regions shown in right-hand close-up images.

### Mitochondrial dysfunction activates the UPR in neurons with high OXPHOS metabolic activity

We next addressed why mitochondrial dysfunction activates the UPR in Imp-expressing central brain B neurons. Central brain B neurons have higher levels of mitochondrial gene expression and are enriched for mitochondrial GO terms compared to central brain A neurons^18^. Consistent with these previous findings, using the expression of 68 mitochondrial oxidative phosphorylation (OXPHOS) genes in the control dataset as an indicator of mitochondrial function using AUCell^18, 20, 27^, central brain B neurons had significantly higher OXPHOS gene expression compared to central brain A neurons, optic lobe neurons and glia (Fig. 4A). Since the UPR was activated in central brain B neurons, we therefore hypothesised that mitochondrial metabolic activity determines the potential to activate the UPR in different neuron types. To test this, we ranked control dataset clusters for OXPHOS gene expression as an indicator of mitochondrial function (Fig. 3B). The relative ranking positions of neuronal and glial cell types in our control dataset was largely consistent with those from single cell RNA-seq analysis of the wild-type *Drosophila* brain^18^. Moreover, in our dataset, head muscle, which is highly dependent on mitochondrial function, had the highest level of OXPHOS gene expression (Fig. 4B). We then first focused on dopaminergic and serotonergic neurons, as these are well characterised genetically tractable central brain neurons and have very low OXPHOS gene expression (Fig. 4B, arrows) and^18^. Pan-neuronal ND-75 KD did not cause ATF4 activation in the large protocerebral anterior medial (PAM) dopaminergic neuron cluster or in posterior dopaminergic neurons (Fig. 4C, D, arrows). Moreover ND-75 KD specifically in dopaminergic and serotonergic neurons, using *TH-Gal4* and *TRH-Gal4* respectively, did not activate ATF4 even at 29°C to maximise Gal4 activity (Fig. 4E, F). Therefore, consistent with their low levels of OXPHOS gene expression and thus relatively low mitochondrial activity, ND-75 KD does not activate the UPR in dopaminergic and serotonergic neurons.

**Fig. 4.**
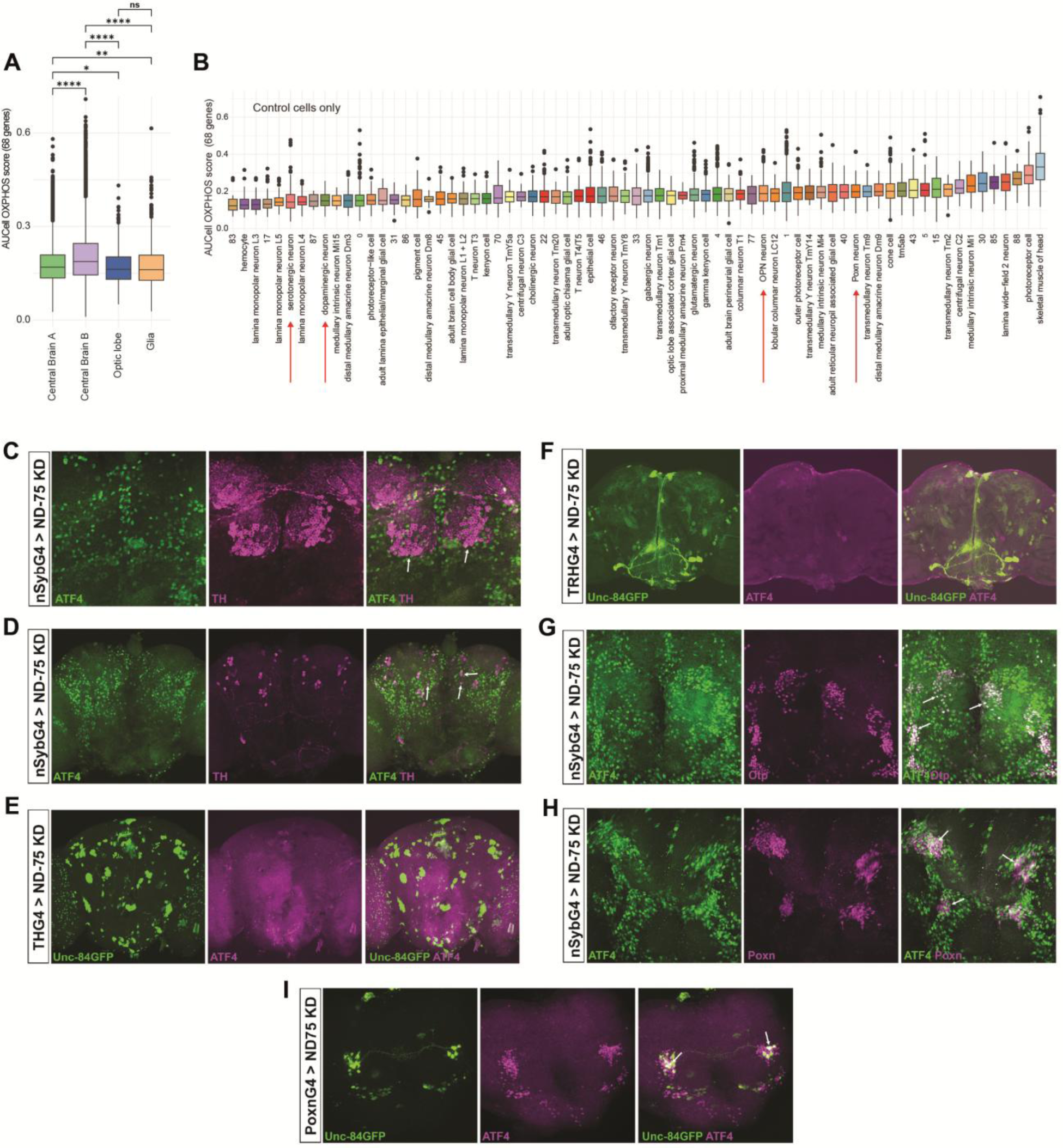
Mitochondrial dysfunction activates the UPR in neurons with high mitochondrial metabolic activity. (A) OXPHOS gene expression in central brain B neurons is significantly higher than central brain A neurons, optic lobe neurons and glia according to Wilcoxon rank-sum tests. (B) Ranking of clusters by expression of 68 OXPHOS genes in control cells only. Arrows highlight dopaminergic, serotonergic, OPN and Poxn clusters. (C, D) Pan-neuronal (*Tub-Gal80^ts^;nSyb-Gal4*) ND-75 KD does not activate ATF4 (green) in dopaminergic PAM neurons (C, arrows) or posterior dopaminergic neurons (D, arrows), marked by expression of tyrosine hydroxylase (TH, magenta). (E) ND-75 KD specifically in dopaminergic neurons using *TH-Gal4* at 29°C does not activate ATF4 expression (magenta). Unc84-GFP expression (green) is shown as a reporter of *TH-Gal4* activity. (F) ND-75 KD specifically in serotonergic neurons using *TRH-Gal4* at 29°C does not activate ATF4 expression (magenta). Unc84-GFP expression (green) is shown as a reporter of *TRH-Gal4* activity. (G) Pan-neuronal (*Tub-Gal80^ts^;nSyb-Gal4*) ND-75 KD activates ATF4 (green) in OPNs, marked by expression of Otp (magenta), arrows. (H) Pan-neuronal (*Tub-Gal80^ts^;nSyb-Gal4*) ND-75 KD activates ATF4 (green) in Poxn neurons, marked by expression of Poxn (magenta), arrows. (I) ND-75 KD specifically in Poxn neurons using *Poxn-Gal4* at 29°C activates ATF4 (magenta), arrows. Unc84-GFP expression (green) is shown as a reporter of *Poxn-Gal4* activity. All brain images are from 1-2 day old flies. ns not significant, *p < 0.05, **P<0.01, ****p < 0.0001.

To further test the hypothesis that mitochondrial activity determines the potential to activate the UPR, we focused on OPNs and Poxn neurons, which scored high for OXPHOS gene expression (Fig. 4B, arrows) and^18^. Orthopedia (Otp) is a homeodomain transcription factor that is expressed in several areas of the adult brain including OPNs^18, 19, 28^. Pan-neuronal ND-75 KD caused ATF4 activation in Otp-expressing OPNs (Fig. 4G, arrows). Poxn is a paired and homeodomain transcription factor expressed in neurons in the central complex^29–31^. Co-staining for Poxn and ATF4 in brains with pan-neuronal ND-75 KD showed ATF4 activation in Poxn-expressing neurons (Fig. 4H, arrows). Moreover, ND-75 KD specifically in GFP-labelled Poxn neurons, using *Poxn-Gal4* at 29°C, caused strong activation of ATF4 (Fig. 4I, arrows). In sum, these data provide robust support for the notion that the UPR is activated by mitochondrial dysfunction in neurons with inherently high mitochondrial activity and not in neurons with low mitochondrial activity.

### Mitochondria-ER contact sites (MERCs) and PERK underlie the susceptibility to activate the UPR in neurons with high mitochondrial metabolic activity

To better understand the mechanism underlying the mitochondrial metabolic activity-dependent activation of the UPR in neurons, we focused on MERCs. Depending on the cell type, a proportion of mitochondria are in close apposition to the ER and have shared functions in lipid transfer, calcium buffering, mitochondrial fission and autophagy^32^. At MERCs, mitochondria and the ER physically interact through tethering complexes and disruption of these tethering complexes activates the UPR^33–35^. We hypothesised that the sensitivity of mitochondrial dysfunction-dependent UPR activation in different neuron types is encoded by MERCs. To test this hypothesis, we used the split-GFP-based contact site sensor (SPLICS) to visualise MERCs (Fig. 5A)^17, 36^. Very surprisingly, pan-neuronal expression using *nSyb-Gal4* in the wild-type brain of SPLICS-long (SPLICS-L), which detects wide (∼40-50 nm) MERCs, resulted in SPLICS-L puncta only in cell bodies in specific central brain regions and a subset of neurons (Fig. 5B). By contrast, pan-neuronal expression of mitochondrially-targeted GFP (mito-GFP), or BiP-sfGFP-HDEL, which labels the ER, resulted in broad uniform localisation throughout the central brain (Fig. 5C, D). Pan-neuronal expression of an independent reporter, SPLICS-short (SPLICS-S), which detects narrow (∼8-10nm) MERCs, gave a very similar expression pattern to SPLICS-L, with puncta only in specific central brain regions and a subset of neurons (Fig. 5E).

**Fig. 5.**
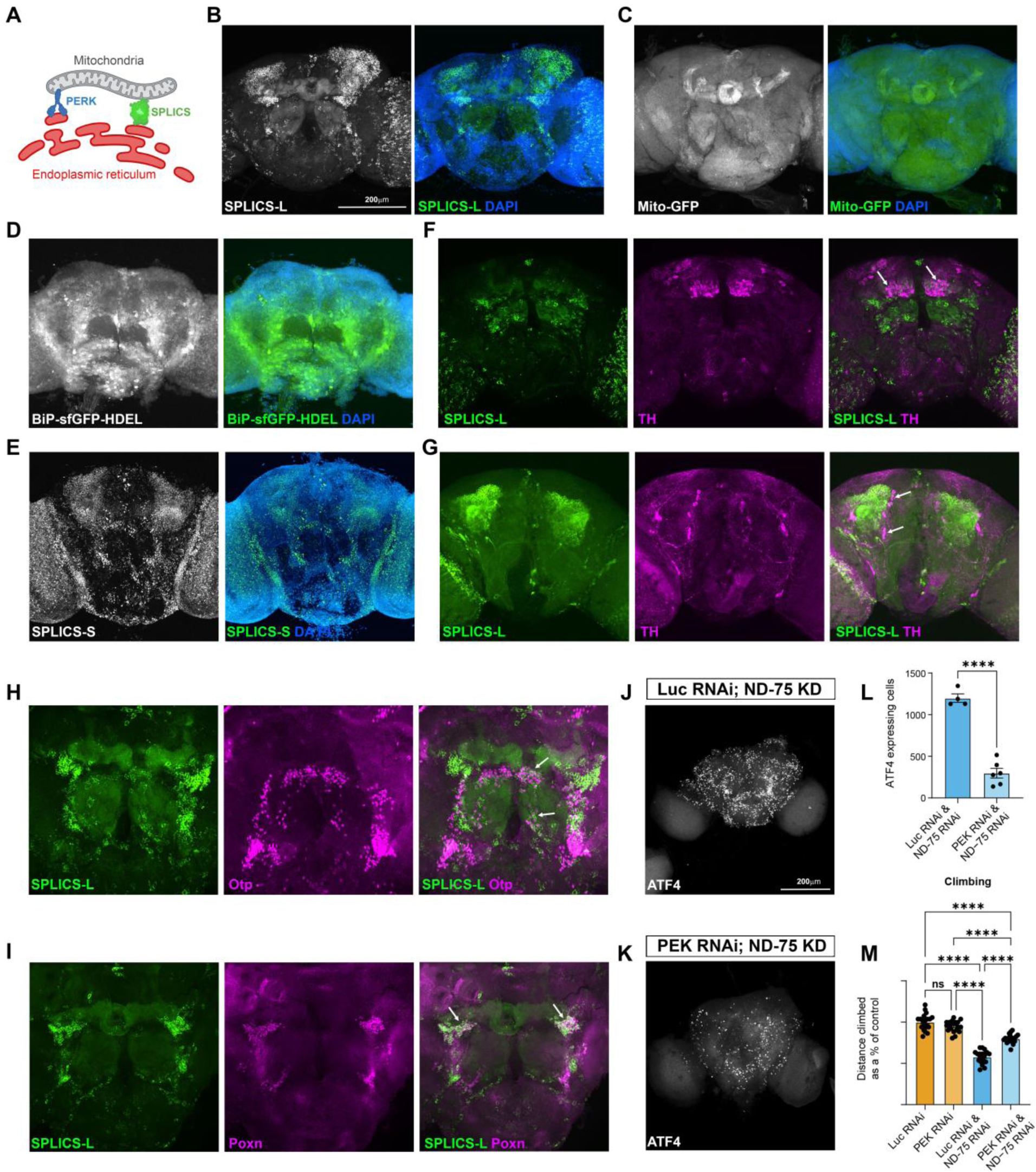
MERCs and PERK underlie neuron-type specific UPR activation. (A) Schematic showing PERK at MERCs and detection of MERCs by SPLICS. (B) Pan-neuronal (*nSyb-Gal4*) expression of SPLICS-L, a reporter of MERCs, is localised to specific brain regions and neurons. (C) Pan-neuronal (*nSyb-Gal4*) expression of mito-GFP, a reporter of mitochondria, is uniformly localised throughout the brain. (D) Pan-neuronal (*nSyb-Gal4*) expression of BiP-sfGFP-HDEL, a reporter of ER, is uniformly localised throughout the brain. (E) Pan-neuronal (*nSyb-Gal4*) expression of SPLICS-S is localised to specific brain regions and neurons. (F, G) With pan-neuronal (*nSyb-Gal4*) expression SPLICS-L (green) is not present in dopaminergic PAM neurons (F, arrows), or posterior dopaminergic neurons (G, arrows), marked by expression of tyrosine hydroxylase (TH, magenta). (H, I) With pan-neuronal (*nSyb-Gal4*) expression high levels of SPLICS (green) are present in OPNs (H, arrows), marked by Otp expression (magenta), and Poxn neurons (I, arrows), marked by Poxn expression (magenta). (J, K) Pan-neuronal (*Tub-Gal80^ts^;nSyb-Gal4*) ND-75 KD with luciferase RNAi (J) or PEK RNAi (K) shows that PEK knockdown reduces the number of neurons activating ATF4. (L) Quantification of ATF4 expressing cells caused by ND-75 KD with luciferase RNAi and PEK RNAi. Luc RNAi;ND-75 KD n=4 brains, PEK RNAi;ND-75 KD n=6 brains. (M) The reduced climbing ability caused by ND-75 KD in Poxn neurons (using *Poxn-Gal4* at 29°C) is partially rescued by knockdown of PEK. Data are represented as mean ± SEM and were analysed using the student’s unpaired t test in (L) and one-way ANOVA in (M). ns not significant, ****p < 0.0001.

Remarkably, even though expressed pan-neuronally, SPLICS-L puncta were not visible in cell bodies of either anterior PAM dopaminergic neurons or posterior dopaminergic neurons (Fig. 5F, G, arrows). By contrast, SPLICS-L puncta were abundant in cell bodies of Otp-expressing OPNs and Poxn-expressing Poxn neurons (Fig. 5H, I, arrows). Thus, under normal conditions, MERCs are abundant in OPNs and Poxn neurons, the neuron types with high mitochondrial activity that activate the UPR upon ND-75 KD, but are undetectable in dopaminergic neurons.

We next expressed mito-GFP or SPLICS-L specifically in neurons with high or low mitochondrial activity using cell type specific Gal4 drivers. High levels of mito-GFP were observed in cell bodies of Poxn neurons (using *Poxn-Gal4*), OPNs (using *GH146-Gal4*), dopaminergic neurons (using *TH-Gal4*) and serotonergic neurons (using *TRH-Gal4*) (Supplementary Fig. S4A-D, arrows). By contrast, expression using the same Gal4 drivers showed that SPLICS-L puncta were clearly visible in cell bodies of Poxn neurons and OPNs (Supplementary Fig. S4E, F, arrows), but almost completely absent in dopaminergic and serotonergic neurons with weak expression visible only in cell bodies of a few posterior dopaminergic neurons and serotonergic neurons in the suboesophageal zone (Supplementary Fig. S4G, H, arrows). These data show that, even when expressed using cell-type specific Gal4 drivers, MERCs were not observed in most dopaminergic and serotonergic neurons. By contrast, MERCs were abundant in Poxn neurons and OPNs. This provides strong support for the hypothesis that inherent quantitative differences in MERCs determine whether neurons activate the UPR in response to mitochondrial dysfunction.

To understand the mechanisms involved, we focused on protein kinase RNA-like ER kinase (PERK). PERK is a component of the UPR that upon ER stress phosphorylates eIF2alpha and activates ATF4 translation^37^. PERK is localised to MERCs and is required to maintain MERCs, where it physically interacts with the mitofusin 2 in the mitochondrial outer membrane (Fig. 5A)^38, 39^. Simultaneous knockdown of PEK, the *Drosophila* homolog of PERK, and ND-75 KD using *Tub-Gal80^ts^;nSyb-Gal4* significantly reduced the number of neurons expressing ATF4 in the brain compared to ND-75 KD alone (Fig. 5J-L). Similarly, PEK knockdown significantly reduced the number of neurons expressing Xrp1-HA in the brain compared to ND-75 KD alone (Supplementary Fig. S5).

To test the functional requirement for PERK and MERCs, we took advantage of the role of Poxn neurons in the regulation of locomotor function^40, 41^. ND-75 KD in Poxn neurons using *Poxn-Gal4* at 29°C caused a strong reduction in climbing ability, which was partially rescued by knockdown of PEK (Fig. 5M). Thus, reducing the expression of PERK, which is localised to MERCs and required for ATF4 activation via the UPR, ameliorates the loss of neuronal function caused by mitochondrial dysfunction in Poxn neurons.

### UPR activation in neurons with high mitochondrial metabolic activity in the human brain

Our findings in *Drosophila* showed that the UPR is activated only in neurons with high mitochondrial metabolic activity (Fig. 6A, B). To determine whether this principle is also true in the human brain, we analysed snRNA-seq data from a recent study of *substantia nigra pars compacta* (SNpc) tissue from 29 PD patients and old-aged controls, with an average age of 79^42^. Ranking all cell types in the SNpc showed that neurons had the highest OXPHOS gene expression (Fig. 6C and^42^). Clustering of the neuronal population identified 6 transcriptionally distinct clusters of cells, termed Neurons0-Neurons5 (Supplementary Fig. S6A)^42^. Within the six clusters, Neurons0 and Neurons3 ranked highest for OXPHOS gene expression (Fig. 6D; Supplementary Fig. S6B). Analysis of the expression of UPR marker genes *HSP90AA1* and *HSPA8* showed that both genes had highest expression in Neurons0 and Neurons3 clusters (Fig. 6E, F; Supplementary Fig. S6C, D). Moreover, expression of the 93 REACTOME UPR gene set^43^ was highest in Neurons0 and Neurons3 clusters (Fig. 6G; Supplementary Fig. S6E). ATF4 regulon activity was also most strongly enriched in Neurons0 and Neurons3 clusters (Fig. 6H; Supplementary Fig. S6F). These data demonstrate that high OXPHOS gene expression correlates with UPR activation in neurons in the SNpc.

**Fig. 6.**
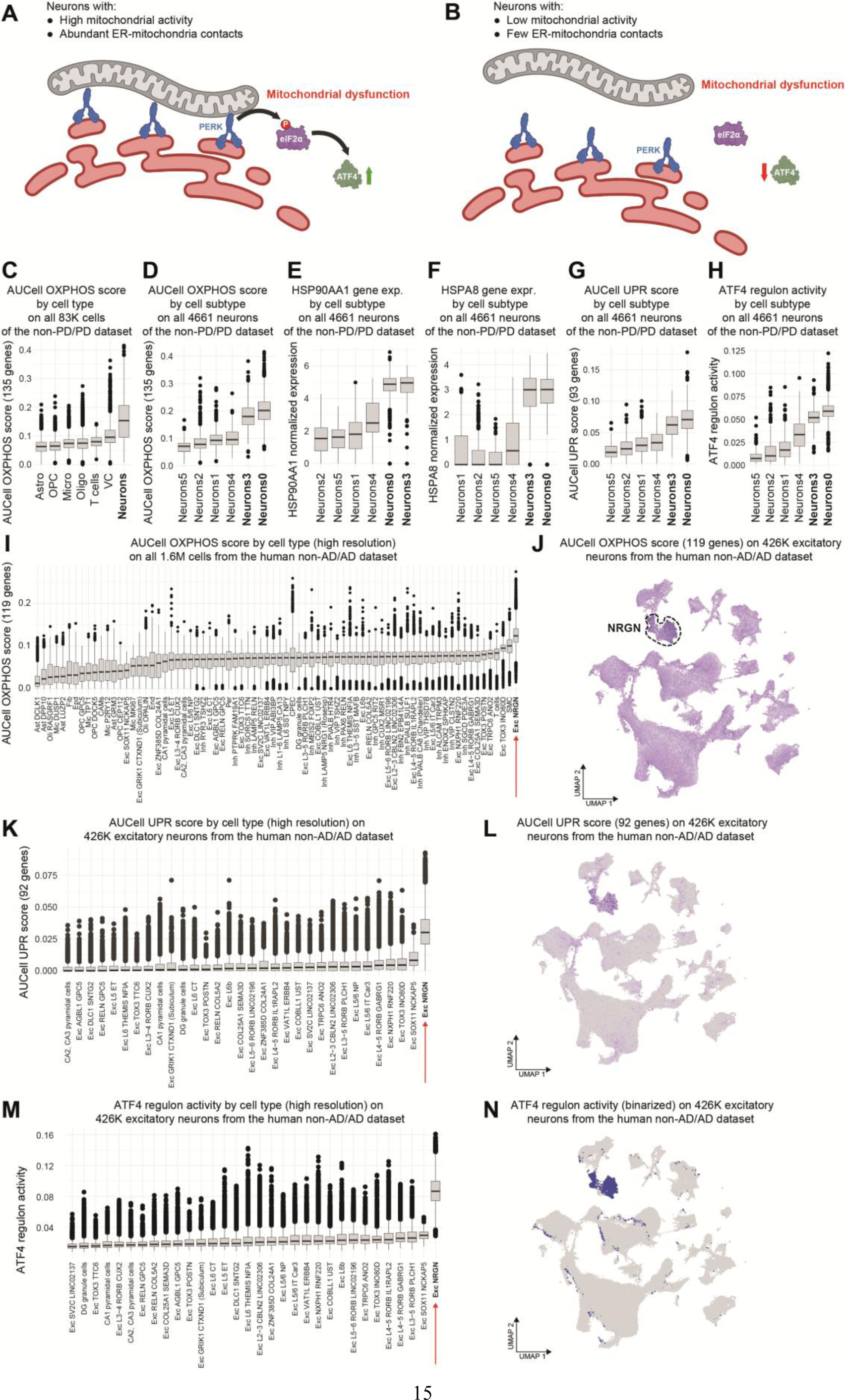
The UPR is activated in neurons with high mitochondrial metabolic activity in the human brain. (A) In neurons with high mitochondrial metabolic activity mitochondrial dysfunction causes ER stress and activates the UPR via PERK. Consequently, mitochondrial dysfunction causes ER stress and activates the UPR via PERK. (B) In neurons with low mitochondrial metabolic activity mitochondrial dysfunction is limited to the organelle and does not activate the UPR. Consequently, mitochondrial dysfunction is limited to the organelle, does not affect the ER and does not activate the UPR. (C) AUCell aggregated score computed on 135 OXPHOS-related genes and ranked for each broad cell-type (83K cells) of the non-PD/PD SNpc dataset^42^ showing highest activity in neurons. (D-H) AUCell aggregated score computed on 135 OXPHOS-related genes (D), *HSP90AA1* normalized gene expression (E), *HSPA8* normalized gene expression (F), AUCell aggregated score computed on 93 UPR-related genes (G), and SCENIC ATF4 regulon activity (H) ranked for each neuron subtype of 4,661 neurons extracted from the non-PD/PD dataset showing highest OXPHOS, UPR and ATF4 activity in Neurons0 and Neurons3 clusters. (I) AUCell aggregated score computed on 119 OXPHOS-related genes ranked for each broad cell-type (1.6M cells) of the non-AD/AD dataset^44^ from 6 brain regions showing highest activity in excitatory NRGN neurons. (J) UMAP representing AUCell aggregated score computed on 119 OXPHOS-related genes for ∼426k excitatory neurons extracted from the non-AD/AD dataset showing highest expression in excitatory NRGN neurons. (K, L) AUCell aggregated score computed on 92 UPR-related genes ranked for each excitatory neuron subtype (K) and UMAP representation (L) of ∼426k excitatory neurons extracted from the non-AD/AD dataset showing highest activity in excitatory NRGN neurons. (M, N) SCENIC ATF4 regulon activity ranked for each excitatory neuron subtype (M) and UMAP representation (N) in the non-AD/AD dataset showing highest activity in excitatory NRGN neurons.

To examine the human brain more broadly, we used snRNA-seq data from a recent study of six brain regions, including the hippocampus and prefrontal cortex, from 48 AD patients and non-AD age-matched individuals ranging in age from 75-95^44^. Ranking of all cell types, or excitatory neurons, showed that excitatory neurogranin (NRGN) neurons had the highest level of OXPHOS gene expression, regardless of age (Fig. 6I, J; Supplementary Fig. S7A, B). Remarkably, excitatory NRGN neurons also ranked by far the highest for UPR gene expression and the UMAP showed that UPR gene expression was highly enriched in NRGN neurons (Fig. 6K, L). Excitatory NRGN neurons also had by far the highest level of ATF4 regulon activity and the UMAP showed that ATF4 regulon activity was almost exclusive to the NRGN cluster (Fig. 6M, N). Interestingly, UPR gene expression and ATF4 regulon activity in NRGN neurons was increased equally in AD patients and aged-matched non-AD control individuals (Supplementary Fig. S7C, D). These data provide further support for the principle that the UPR is activated in neurons with high mitochondrial metabolic activity in the human brain.

## Discussion

The cell-type specificity of stress signaling pathways in the brain is largely uncharted Here, we revealed how specific types of neurons in the *Drosophila* brain respond to mitochondrial dysfunction by activating the UPR and transforming their transcriptional identity. The UPR was activated primarily in central brain B neurons that had high mitochondrial activity. We also discovered a previously unknown remarkable difference in the abundance of MERCs throughout the brain, which are almost undetectable in neurons with low mitochondrial activity.

Mitochondrial metabolic activity and abundance of MERCs in neurons encodes the potential to activate the UPR and requires the ER kinase PERK. Finally, the UPR was activated in the human brain in neurons with the highest mitochondrial activity. Overall, this study reveals the fundamental dependence of the UPR on mitochondrial metabolic activity in the brain.

Transcriptional reprogramming is an essential mechanism in nervous system development^45–47^. Using snRNA-seq to dissect the transcriptome in single cells in the ND-75 KD brain, we showed that transcriptional identity was largely preserved in neurons with mitochondrial dysfunction. However, unlike most cell types in the brain of ND-75 KD flies, clusters 2 and 23 consisted almost exclusively of ND-75 KD cells. Regulon analysis also showed that genes regulated by crc/ATF4, a core component of the UPR, were enriched most strongly in cluster 23. The UPR is primarily a stress response pathway that promotes either an adaptive response to restore cellular homeostasis or apoptotic cell death ^8^. However, the UPR also has physiological roles in promoting the differentiation of B cells in the immune system and differentiation of epidermal keratinocytes^48, 49^. We performed ND-75 KD in differentiated postmitotic neurons and, although cell fate is usually stable at this stage, mitochondrial dysfunction reprogrammed the transcriptional identity of a specific subset of neurons in the central brain. Presumably, the common transcriptional response to mitochondrial dysfunction overwhelmed the normal transcriptional signature in these neurons. Activation of the UPR contributed to loss of neuronal function, as knockdown of PEK improved the motor function of flies with ND-75 KD in Poxn neurons. Chronic activation of mitochondrial stress signaling via the UPR therefore contributes to neuronal dysfunction.

Mitochondrial dysfunction has been shown to activate the UPR in multiple cell types and disease contexts^50–57^. Cell-type specificity of stress signaling pathways is not evident from experiments in cultured cell lines, or using tissue homogenate, but can be revealed using single cell and single nuclear transcriptomic approaches. The novel clusters we identified through snRNA-seq of the ND-75 KD brain could potentially derive from a single cell type that is highly sensitive to mitochondrial dysfunction, or from multiple cell types that converge to a common unique transcriptional identity. Our data indicate that clusters 2 and 23 derive from the reprogramming of multiple neuron types that are highly sensitive to mitochondrial dysfunction. Mitochondrial activity was previously identified as a key variable distinguishing different types of neurons in the *Drosophila* brain^18^. Neurons in the central brain are either central brain B neurons that express Imp and have higher mitochondrial activity, or central brain A neurons that express Pros/Dati and have lower mitochondrial activity. We found that mitochondrial dysfunction activates the UPR primarily in Imp-expressing, more mitochondrially active, central brain B neurons.

In the context of the nervous system, neurodegenerative diseases preferentially affect specific cell types and brain regions, emphasising the importance of understanding the cell-type specific mechanisms underlying the response to stress in the brain. Our work clearly revealed that pan-neuronal mitochondrial dysfunction activates the UPR in a specific subset of neurons in the central brain. This specificity is encoded by the underlying mitochondrial metabolic activity and abundance of MERCs in different neuron types. Our data suggest a model where neurons with high mitochondrial metabolic activity require close association with the ER, through abundant MERCs (Fig. 6A). However, if mitochondria are dysfunctional, this close association results in ER stress and activation of the UPR, which coordinates a transcriptional response to adapt to ER stress (Fig. 6A). By contrast, in neurons with low mitochondrial metabolic activity and few MERCs, mitochondrial dysfunction is contained within the organelle and does not cause ER stress nor activate the UPR (Fig. 6B). Thus, cellular stress signaling specificity in the brain derives from each neuron’s inherent mitochondrial activity and the proximity of mitochondria and the ER, which encodes its potential to activate the UPR.

If our model is generalisable, the principle that mitochondrial metabolic activity determines the types of neurons that activate the UPR should be conserved across species. Applying the same computational analysis used in *Drosophila* to snRNA-seq data from human brain tissue showed that different neuron types exhibit varying levels of OXPHOS gene expression. Consistent with our model, neurons in the SNpc with high OXPHOS gene expression showed evidence of UPR activation. Moreover, excitatory NRGN neurons had the highest OXPHOS gene expression and showed the strongest UPR activation in the brain of aged individuals. NRGN regulates synaptic signaling, interacts with calmodulin and is required for calcium/CaM kinase II-dependent long term potentiation^58^. NRGN neurons may therefore be highly sensitive to mitochondria-ER calcium buffering. Overall, our findings show that the relationship between mitochondrial metabolic activity and UPR activation in neurons is conserved between *Drosophila* and humans. The inherent oxidative metabolic activity of neurons underlying their potential to activate the UPR may therefore be a general principle.

## Methods

### Fly strains and growth conditions

Flies were maintained on standard food (per litre: 6.4 g Agar (Fisher), 64 g glucose (Sigma), 16 g ground yellow corn and 80 g Brewer’s yeast (MP Biomed Europe), 3 ml propionic acid (Fisher), 1.8 g methyl 4-hydroxybenzoate (Sigma), 16 ml ethanol (Sigma)) at 25°C in a 12 hour light/dark cycle unless stated otherwise. Genotypes of the *Drosophila* lines used are described in Supplementary Table S1. ND-75, PEK and luciferase RNAi lines were from the TRiP collection^59^ and obtained from the Bloomington Drosophila Stock Center.

### Behavioural analysis

For experiments using ND-75 KD with *nSyb-Gal4* vials were placed at 45° during eclosion to prevent flies from becoming stuck in the food. Climbing assays were performed 1-3 hours after daily illumination (between 8:00am and 11:00am). Adult male flies of the desired genotype were selected on the day of eclosion under CO_2_ anaesthesia. The following morning, individual 1-2 day old flies were transferred into a 10ml serological pipette (Falcon) using a mouth aspirator. Flies were tapped to the base of the pipette, the height climbed by each fly in three continuous 10 second climbs measured and the mean calculated. Any runs where the fly paused midway through the climb or moved horizontally were excluded.

### Immunofluorescence and imaging

A mix of male and female 1-2 day old flies were used except where stated otherwise. Flies were briefly anaesthetised using CO_2_ and washed in ethanol before being transferred to a Sylgard dish. Flies were pinned down upside down (ventral side up) using insect pins (AA2 0.10mm, Watkins and Doncaster) through the thorax. Pinned flies were then moved into pools of PBS on the dish. Adult wings and the first set of legs were removed, the brain dissected by first pushing through the antenna with tweezers (Dumont), removing the eyes, then the rest of the head cuticle. Adult brains were placed in 150µl of 4% formaldehyde (Agar Scientific)/PBS in a 96-well-plate on ice using tweezers, repeated for 10-15 brains per genotype, then incubated on ice for 20 minutes to fix the tissue. Fixative was removed and brains were then washed twice in PBS/0.1% Triton X100 (Sigma Aldrich) (PBST). Brains were then blocked by incubating in 150µl PBST + 10% normal goat serum (Thermo Fisher) for 60 minutes on a platform rocker. The tissue was incubated overnight at 4°C with primary antibody diluted in PBST + 10% NGS on a platform rocker in a small volume (100-200µl) in a 96-well-plate. The primary antibody was then removed and brains washed in 150µl PBST 4x for 10 minutes at room temperature on a platform rocker. The tissue was then incubated with 150µl secondary antibody diluted 1/1000 in PBST + 10% NGS at room temperature on platform rocker for 1.5 hours with the 96-well-plate was wrapped in foil, followed by 4x washes in PBST for 10 minutes with the 96-well-plate wrapped in foil on platform rocker. PBST was then replaced with PBS. A P200 tip with the end cut off was used to pipette the brains onto a Superfrost microscope slide (Thermo scientific). Excess PBS was dabbed away using paper tissue and 15µl of Vectashield mounting media (Vector laboratories) added followed by a 1.5 thickness, 22 x 22mm coverslip (VWR) gently on top of the slide. The cover slip was held in place using nail varnish. Slides were stored at 4°C in the dark. A Nikon AX inverted confocal microscope was used for imaging of brains. Z stacks were taken using either an Apo 20x lens (NA=0.75) or an Apo 60x oil immersion lens (NA=1.49). All images were taken at 1024×1024 pixels in size. Brain images were quantified using 3D Z-stack images or 2D projection images and regions of interest analysed in Volocity (Perkin Elmer) using the Measurement tool.

The following primary antibodies were used. Rat anti-ATF4 (1/300, preabsorbed against dissected larval tissues ^4^); mouse anti-TH (1/50, GmbH ABIN617906); rabbit anti-Poxn (1/200, a gift from Frank Hirth, ^60^), mouse anti-NC82 (1/200, DSHB); guinea pig anti-Otp (1/4000, a gift from Uwe Walldorf, ^28^); mouse anti-HA (1/100, Cell Signalling 2367); mouse anti-Pros (1/2, DSHB). Secondary antibodies were all used at 1/1000 and were goat anti-mouse Alexa Fluor 488 (Thermo Fisher A11001), got anti-rat Alexa Fluor 555 (Thermo Fisher A21434), goat anti-rabbit Alexa Fluor 546 (Invitrogen A11035) and goat anti-rabbit Alexa Fluor 633 (Invitrogen A21071).

### Drosophila snRNA-seq

*Tub-Gal80^ts^;nSyb-Gal4* virgin females were crossed to either UAS-Unc84-GFP males (control) or UAS-Unc84-GFP;ND-75 RNAi males (ND-75 KD) and progeny incubated at 25°C throughout development then transferred to 29°C immediately after eclosion for 24 hours to enhance Gal4 activity. Male flies were snap frozen and heads collected. 60 fly heads per genotype were collected and stored at −80°C in 1.5ml tubes.

#### Nuclei isolation

First, 200µl of homogenisation buffer (250mM sucrose (Fisher scientific), 10mM Tris pH 8.0 (Sigma Aldrich), 25mM KCL (Applichem), 5mM MgCl_2_ (Sigma Aldrich), 0.1% Triton x 100 (Sigma Aldrich), 0.1% DTT (NEB), 1x protease inhibitor (Thermo scientific), 0.5% Rnasin ribonuclease inhibitor (Promega), nuclease free water (Invitrogen)) was added to a 1.5ml Eppendorf with 60 flash frozen heads. A motorised pestle (Grainger) was used to grind samples for 2 minutes on ice. A further 800µl homogenisation buffer was added, and the sample was transferred to a 1ml glass dounce pestle (DWK Life Sciences). 60 strokes were applied in the tight pestle. The sample was then filtered through a 35µm cell strainer into a 5ml FACs tube (Corning). Next, the sample was transferred into a 1.5ml Eppendorf and centrifuged for 10 minutes at 1,000g. The supernatant was discarded, and the sample was re-suspended in 500µl resuspension buffer (0.5% BSA (Sigma Aldrich), 0.5% Rnasin ribonuclease inhibitor (Promega), 1x PBS (Gibco)). The sample was then filtered through a 40µm Flowmi cell strainer (Bel-Art) into a 5ml FACs tube. Prior to sorting, 20µl of unstained nuclei were added to 200µl resuspension buffer as an unstained control for setting up the FACs gate. Hoechst-33342 (1:1000, (Thermo Fisher)) was added to the remaining nuclei and incubated for 5 minutes. The sample was then sorted for Hoechst+ nuclei into a collection tube. After FACs sorting, the collection tube was centrifuged for 10 minutes at 1,000g. The supernatant was then discarded and the pellet re-suspended in 45.2µl resuspension buffer. This protocol was repeated for each genotype.

#### GEM generation and barcoding

The Chromium Next GEM Single Cell 3’ Kit v3.1 was used for GEM generation and barcoding. 31.9µl Master Mix (18.8µl RT Reagent B, 2.4µl Template Switch Oligo, 2.0µl Reducing Agent B, 8.7µl RT Enzyme C) was added into each tube of a 8-tube PCR strip on ice. The Chromium Next Gem Chip was then removed from the sealed bag, and the chip used within 24 hours of opening. To prepare the Master Mix + Cell suspension 26.7µl of nuclease free water was added to the Master Mix and added to 16.5µl of cell suspension stock (1000 cells/µl) so the total volume was 75µl in each tube. Samples were loaded into row 1. Gently, 70µl of Master Mix + Cell suspension was pipetted into the bottom centre of each well.

The Gel Bead strips were vortexed for 30 seconds and centrifuged for 5 seconds. Samples were loaded into row 2. Slowly, 50µl Gel Beads was aspirated and dispensed into each well. Samples were loaded into row 3.45µl Partitioning Oil was dispensed into wells. The GEM Gasket was attached, aligned notch with top hand corner and ensured the gasket holes are aligned with the wells. The Chromium Controller was run for 18 minutes. After, GEMs were transferred. The gasket was discarded and 100µl GEMs were slowly aspirated, the GEMs were dispensed into a tube strip on ice. The GEMs were incubated in a thermocycler (Bio-Rad) set at: (53°C for 45 minutes, 85°C for 5 mins and 4°C hold). Stored at 4°C for up to 72 hours.

#### Post GEM-RT cleanup and cDNA amplification

The Chromium Next GEM Single Cell 3’ Kit v3.1 was used for Post GEM-RT cleanup and cDNA amplification. The post GEM-RT clean-up was performed with Dynabeads. 125µl recovery agent was added to each sample at room temperature and left for 2 minutes. Slowly, 125µl recovery agent/partitioning oil (pink) was removed and discarded from the bottom of the tube. Next, 200µl of Dynabeads Cleanup Mix (182µl Cleanup Buffer, 8µl Dynabeads MyOne SILANE, 5µl Reducing Agent B, 5µl Nuclease-free water) was added to each sample. The samples were incubated for 10 minutes at room temperature. The Elution Solution I (98µl Buffer EB, 1µl 10% Tween 20, 1µl Reducing Agent B) was prepared. The samples were placed on a 10x Magnetic Separator-High position until the solution cleared. The supernatant was removed. 300µl 80% ethanol was added to the pellet while on the magnet. The samples were left to wait for 30 seconds and ethanol was removed. 200µl 80% ethanol was added to the pellet and left to wait for 30 seconds. The ethanol was removed and left to air dry for 1 minute. The magnet was removed and 35.5µl Elution Solution I was added. The mix was pipetted and incubated for 2 minutes at room temperature. Samples were placed on magnet-low until the solution cleared. 35µl sample was transferred to a new tube strip. The cDNA Amplification Mix (50µl Amp Mix, 15µl cDNA primers) was then prepared on ice. 65µl cDNA Amplification Mix was added to 35µl sample. The mix was pipetted 15x and centrifuged briefly. Next, the mix was incubated in a thermal cycler set to: 98°C for 3 minutes, 98°C for 15 seconds, 63°C for 20 seconds, 72°C for 1 minute, for 11 cycles, then 72°C for 1 minute and 4°C to hold. Stored at 4°C for up to 72 hours.

The samples were then vortexed to resuspend the SPRIselect reagent. 60µl SPRIselect reagent was added to each sample and pipetted mix 15x. Samples were incubated for 5 minutes at room temperature. Then, samples were placed on the magnet-high until the solution cleared. The supernatant was removed. 200µl 80% ethanol was added to the pellet and left for 30 seconds, before removing the ethanol. This was repeated with another 2 ethanol washes. Samples were centrifuged briefly and placed on magnet-low. Any remaining ethanol was removed and air dried for 2 minutes. Samples were removed from the magnet and added 40.5µl Buffer EB, pipetted mix 15x. The mix was incubated for 2 minutes at room temperature. Then, the tube strip was placed on magnet-high until the solution cleared. 40µl sample was transferred to a new tube strip. Tube strip was stored at 4°C for up to 72 hours. 1µl of sample was run on an Agilent Bioanalyzer High Sensitivity chip (Agilent) to determine cDNA total yield in ng.

#### Gene Expression Dual Index Library Construction

The library construction kit was used for 3’ Gene Expression Dual Index Library Construction. The Fragmentation Mix was prepared on ice (5µl Fragmentation Buffer, 10µl Fragmentation Enzyme). 10µl of purified cDNA sample was transferred from the Pellet Cleanup to a tube strip. 25µl Buffer EB was added to each sample. 15µl Fragmentation Mix was added to each sample. The mix was pipetted 15x and centrifuged briefly. The sample was transferred into a thermal cycler with the following incubation protocol: 32°C for 5 minutes, 65°C for 30 minutes and 4°C hold.

30µl SPRIselect reagent was then added to each sample. The mix was pipetted 15x. Samples were incubated for 5 minutes at room temperature. Then, samples were placed on the magnet-high until the solution cleared. 75µl supernatant was transferred to a new tube strip. 10µl SPRIselect reagent was added to each transferred supernatant. The mix was pipetted 15x. Samples were incubated for 5 minutes at room temperature. The samples were placed on the magnet-high until the solution cleared. The supernatant was removed. 125µl 80% ethanol was added to the pellet and then removed. Samples were washed another 2 times in ethanol. Then, centrifuged briefly and placed on magnet-low until the solution cleared. The remaining ethanol was removed. Samples were removed from the magnet and added 50.5µl Buffer EB to each sample. The mix was pipetted 15x and incubated for 2 minutes at room temperature. Samples were placed on magnet-high until the solution cleared. Next, 50µl of sample was added to a new tube strip.

The Adaptor Ligation Mix (20µl Ligation Buffer, 10µl DNA Ligase, 20µl Adaptor Oligos) was prepared and centrifuged briefly. 50µl Adaptor Ligation Mix was then added to 50µl sample, mixed 15x and centrifuged briefly. Samples were incubated in a thermal cycler at 20°C for 15 minutes and 4°C hold. 80µl of SPRIselect reagent was then added to each sample, pipetted 15x and incubated for 5 minutes at room temperature, then placed on magnet-high until the solution cleared. The supernatant was then removed and 200µl 80% ethanol added to the pellet. The ethanol was then removed, 2 ethanol washes performed before being placed on magnet-low. Any remaining ethanol was removed and the pellet was left to air dry for 2 minutes. The samples were then removed from the magnet, 35µl Buffer EB added and then mixed by pipetting 15x. The samples were then incubated for 2 minutes at room temperature, placed on magnet-low until the solution cleared, then 30µl sample was transferred to a new tube strip.

Next, 50µl of Amp Mix was then added to 30µl sample. 20µl of an individual Dual Index TT Set A was added to each sample and the well ID used was recorded. The mix was pipetted 15x and centrifuged briefly. Samples were incubated in a thermal cycler set to the protocol: 98°C for 45 seconds, 98°C for 20 seconds, 54°C for 30 seconds, 72°C for 20 seconds (8-10 cycles with 500-1,000ng cDNA), 72°C for 1 minute and 4°C hold. Samples were stored at 4°C for up to 72 hours.

60µl SPRIselect reagent was then added to each sample and pipetted to mix 15x. Samples were incubated for 5 minutes at room temperature. Then samples were placed on the magnet-high until solution cleared. 150µl supernatant was transferred to a new tube strip. 20µl SPRIselect was added to each transferred supernatant. The mix was pipetted 15x and incubated for 5 minutes at room temperature. Samples were placed on the magnet-high until the solution cleared. 165µl supernatant was removed. With the tube still in the magnet, 200µl 80% ethanol was added to the pellet. The ethanol was then removed. This was repeated for another 2 washes of ethanol. Samples were centrifuged briefly and placed on magnet-low. Then, samples were removed from the magnet and 35.5µl Buffer EB was added. The mix was pipetted 15x and incubated for 2 minutes at room temperature. Samples were placed on magnet-low until the solution cleared. 35µl of the sample was transferred to a new tube strip. Samples were stored at 4°C for 72 hours or at −20°C for longer term storage. Lastly, 1µl sample was run on an Agilent Bioanalyzer High Sensitivity chip (Agilent) to determine average fragment size.

#### Illumina sequencing

Paired-end, dual indexing sequencing was used. Sequencing reads with recommended number of cycles as follows: Read 1 for 28 cycles, i7 index for 10 cycles, i5 index for 10 cycles, Read 2 for 90 cycles. Loaded 270pM sample to HiSeq 4000 instrument generated FASTQ files ready for snRNAseq analysis.

#### snRNA-seq preprocessing

FASTQ files comprising two 10x libraries (Pan_neuro_control and Pan_neuro_ND75KD) were each processed (alignment, barcode assignment and UMI counting) with the “cellranger count” (version 7.1.0) pipeline. It generated two UMI count matrices of 18,105 genes (from the dm6 genome assembly) by 11,790, and 11,518 cells in Pan_neuro_control and Pan_neuro_ND75KD, respectively.

The .h5 files generated by Cell Ranger in the previous step were loaded in R (v4.3.1) and a Seurat object was created for each individual 10x single-nuclei library using the standard Seurat (v4.4.0) pipeline^61^. Both datasets were filtered independently to remove cells with low number of UMIs, low number of detected genes or high number of mitochondrial reads. See detailed code at https://github.com/DeplanckeLab/Neuro_Droso_ND75KD.

#### Integration of transcriptomic data

The two previously generated Seurat objects were integrated using Harmony (v.1.2.3). A lenient integration method and an “over-corrected” integration were used to validate the independence of the two new ND-75 KD specific clusters. Then, the standard Seurat pipeline was run, i.e. normalization, clustering, and differential expression analyses were performed to identify distinct cell populations and their transcriptional profiles. In particular, we used the Jackstraw algorithm to select the optimal number of principal components (100 in this case) for UMAP and clustering. The 90 annotated clusters found at resolution 4 were manually assigned to specific cell types using known cell type markers from ^18^ and the Fly Cell Atlas^19^, and the visualization and collaborative capacity of the ASAP web portal^62^. See detailed code at https://github.com/DeplanckeLab/Neuro_Droso_ND75KD.

### Parkinson’s and Alzheimer’s human brain atlases

Two external scRNA-seq human atlases were downloaded and processed along with our own dataset, for comparison and validation purposes. The Parkinson’s disease dataset (noted PD) from Martirosyan et al.^42^ was downloaded from GEO with accession number <u>GSE243639.</u> Specifically, we reconstructed a Seurat object from the raw count matrix and added the UMAP and clinical data from the associated metadata files. The Alzheimer’s disease dataset (noted AD) from Mathys et al.^44^ was downloaded from the Synapse portal with accession number <u>syn52293442</u>. We specifically used the Gene Expression (snRNAseq – 10x) processed, multi-region h5ad dataset, that we transformed into a Seurat object, and mapped clinical data using patient ids retrieved from https://ad-multi-region.cells.ucsc.edu/.

### Regulon calculation using pySCENIC

The pySCENIC^20^ pipeline in Python was run from a Docker container (aertslab/pyscenic:0.12.1). Of note, for *drosophila melanogaster*, we did not use the list of TFs provided along with the pySCENIC pipeline, but an updated list that we downloaded from Flybase, and which is available on the <u>GitHub repo of this manuscript</u>, along with all necessary code for reproducibility. In short, we applied the default SCENIC pipeline: 1) Gene regulatory network inference using GRNboost2 with default parameters, 2) Regulon prediction using cisTarget with the following parameters: “rho_mask_dropouts=True, keep_only_activating=True, weighted_recovery=False, rank_threshold=1500, nes_threshold=3, motif_similarity_fdr=0.001, auc_threshold=0.05 and filter_for_annotation=False”, and 3) Enrichment of the regulons using AUCell with default parameters. This provided a numeric regulon enrichment matrix of 575 regulons and 23179 cells. Finally, we applied the binarize() function to obtain a binarized matrix of the same size, specifying active regulons (1) and inactive regulons (0) for each cell. Both regulon outputs were stored as .tsv files to be further imported into the Seurat object. A “regulon UMAP” was then calculated in Seurat, using the regulon matrix as embeddings (instead of the PCA or the Harmony embeddings). We proceeded similarly for calculating the ATF4 regulon activity of the two human datasets (AD and PD)^42, 44^ using the hg38 human TF list from <u>cistarget</u>.

### OXPHOS scoring using AUCell

To create an OXPHOS gene expression score, we used the AUCell tool in the pySCENIC pipeline^20^. We first generated a LOOM file from a Seurat object (whether our integrated dataset, or other objects made from publicly available datasets) which was further read by the pySCENIC pipeline. A Jupyter Notebook script running on a pySCENIC Docker was used to read both this LOOM file and the OXPHOS/UPR gene lists. Gene lists were acquired differently depending on the species: we used the 68 OXPHOS genes from^27^ for *drosophila melanogaster* and the 119 OXPHOS genes from the KEGG database (hsa00190) for *homo sapiens.* Similarly, for the UPR pathway, we used the 93 UPR genes coming from the <u>REACTOME_UNFOLDED_PROTEIN_RESPONSE_UPR</u> pathway, downloaded from the GSEA/MSigDB database^43, 63^. pySCENIC pipeline generates a tab-separated cell/score mapping TSV file, which is then incorporated later as a new metadata in the Seurat object.

### Neuron type categorization

We categorized all cells into 4 different class: Central Brain A, Central Brain B, Optic Lobe, and Glia cells, using a similar method than^18^. Based on the binarized regulon activity matrix, we first assigned the “Optic Lobe” cells as the ones having the scro regulon activated. Then, we repeated this, and assigned the “Glia” cells as the ones with the repo regulon activated. Finally, we assigned the “Central Brain A” cells as the ones with both the dati and pros regulons activated and the “Central Brain B” as the rest. Of note, after assignment, we find that Imp is significantly more expressed in Central Brain B (average normalized expression = 0.742) compared to Central Brain A cells (average normalized expression = 0.574) using a Wilcoxon rank-sum test (p-value = 1.787E-32).

### Final Seurat object and export to the ASAP web portal

The metadata was added with SCENIC regulon data, AUCell enrichment of OXPHOS genes and manual annotation of clusters to the Seurat object. The final Seurat object was then transformed into a .loom file and uploaded to the ASAP web portal (asap.epfl.ch.) ^62^.

### Statistical analyses

Boxplots representations of the snRNA-seq data were made with ggplot2 in R, following standard Tukey-style. Horizontal line = median (or Q2), bottom of box = Q1 (25% quantile), top of the box = Q3 (75% quantile), lines are Q1 - 1.5 IQR (InterQuartile Range) and Q3 + 1.5 IQR. Wilcoxon rank-sum test was used for statistical analysis. Other data were expressed as mean +-S.E.M and analysed using Prism 9 (GraphPad). A student’s unpaired two-way t-test was used for pairwise comparisons of continuous data. An F-test was used to test for any unequal variances, and if significant, Welch’s correction was applied. A one-way ANOVA with Tukey’s post-hoc test was used for multiple comparisons of continuous data. P<0.05 considered statistically significant; * *p*≤0.05, \*\**p*≤0.01, *** *p*≤0.001, **** *p*≤0.0001, n.s. non-significant.

## Data availability

An interactive visualization of the dataset will be made available through the ASAP portal upon acceptance of the manuscript. Raw sequencing data, count matrices, Seurat objects, Loom and h5ad files were deposited on ArrayExpress under accession number https://www.ebi.ac.uk/biostudies/arrayexpress/studies/E-MTAB-14766?key=93078d92-c234-483e-9ee1-3847fe38cd06. Bash, R and Python scripts used for preprocessing and downstream analysis, as well as full scripts for figure panel reproducibility are available on GitHub at https://github.com/DeplanckeLab/Neuro_Droso_ND75KD.

## Acknowledgements

We are grateful to Frank Hirth, Nick Baker, Alessio Vagnoni and Hongjie Li for fly stocks, Uwe Walldorf and Frank Hirth for antibodies, Lucy Granat and Navdeep Chandel for helpful comments on the manuscript. Stocks obtained from the Bloomington *Drosophila* Stock Center (NIH P40OD018537) were used in this study. We thank the Wohl Cellular Imaging Centre at King’s College London for help with light microscopy.

## Funding

The work was funded by grants from the NC3Rs (NC/V001884/1) and the MRC (MR/V013130/1) to JMB and Swiss National Science Advanced Grant (#TMAG-3_209335), Swiss Institute of Bioinformatics ASAP Resource, and EPFL School of Life Sciences Institutional funding to BD.

## Author contributions

Conceptualization: BD, JMB

snRNA-seq: ELH, RF

Computational analysis: VG

In vivo data validation: ELH, JMB

Writing: JMB

Review and Editing: All authors

Supervision: BD, JMB

Funding Acquisition: BD, JMB

## Competing interests

Authors declare that they have no competing interests

## Notes

### Competing Interest Statement

The authors have declared no competing interest.

