## Supplemental figures S1-S7, Supplemental table S1 for "The cellular basis of mitochondrial stress signaling in the brain"

### Supplementary Figures

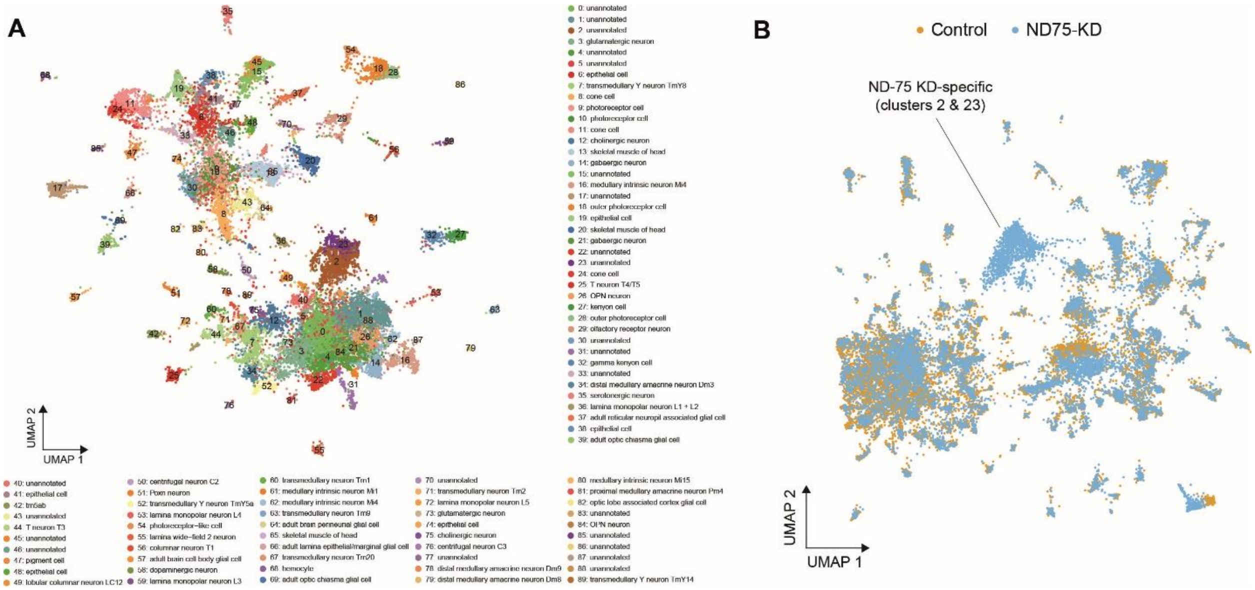


*Fig. S1. ND-75 KD reprogrammes the transcriptional identity of a specific subset of neurons*. (A) UMAP listing and colour coding of all annotated and unannotated cell clusters in the Harmony-integrated snRNA-seq dataset. (B) UMAP without Harmony integration showing control (orange) and ND-75 KD (blue) cell identities. Novel clusters 2 and 23 consist almost exclusively of ND-75 KD cells.

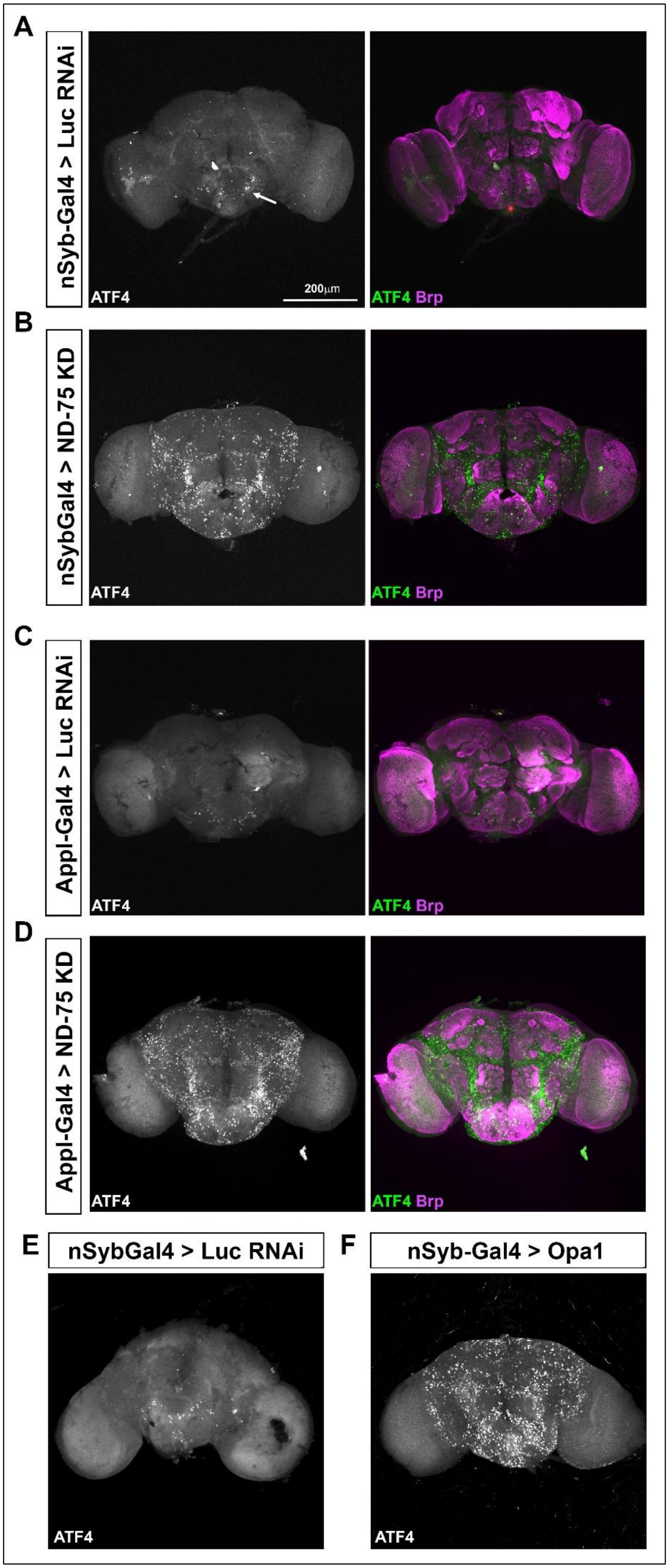

A
nSyb-Gal4 > Luc RNAi
ATF4
200μm
ATF4 Brp
B
nSybGal4 > ND-75 KD
ATF4
ATF4 Brp
C
Appl-Gal4 > Luc RNAi
ATF4
ATF4 Brp
D
Appl-Gal4 > ND-75 KD
ATF4
ATF4 Brp
E
nSybGal4 > Luc RNAi
ATF4
F
nSyb-Gal4 > Opa1
ATF4

*Fig. S2. Mitochondrial dysfunction activates ATF4 in a specific subset of neurons in the central brain.* (A, B) ATF4 (white on left, green on right) is only expressed in a small number of cells in the suboesophageal ganglion (arrow) in *Tub-Gal80$^{ts}$;nSyb-Gal4*-driven luciferase RNAi control brain (A), but is strongly activated throughout the central brain in a specific subset of neurons with pan-neuronal (*Tub-Gal80$^{ts}$;nSyb-Gal4*) ND-75 KD at 25ºC (B). Brp staining (magenta) shows overall brain structure and anatomical regions. (C) Pan-neuronal expression of luciferase RNAi with *Appl-Gal4;Tub-Gal80$^{ts}$* stained for ATF4 (white on left, green on right) and Brp (magenta). (D) ND-75 KD with *Appl-Gal4;Tub-Gal80$^{ts}$* at 25ºC causes strong ATF4 (white on left, green on right) activation throughout the central brain in a specific subset of neurons in a very similar pattern to ND-75 KD with *Tub-Gal80$^{ts}$;nSyb-Gal4*. Brp staining (magenta) shows overall brain structure and anatomical regions. (E, F) Pan-neuronal (*Tub-Gal80$^{ts}$;nSyb-Gal4*) Opa1 overexpression at 29ºC in the late pupal brain causes ATF4 activation throughout the central brain in a specific subset of neurons in a very similar pattern to ND-75 KD.

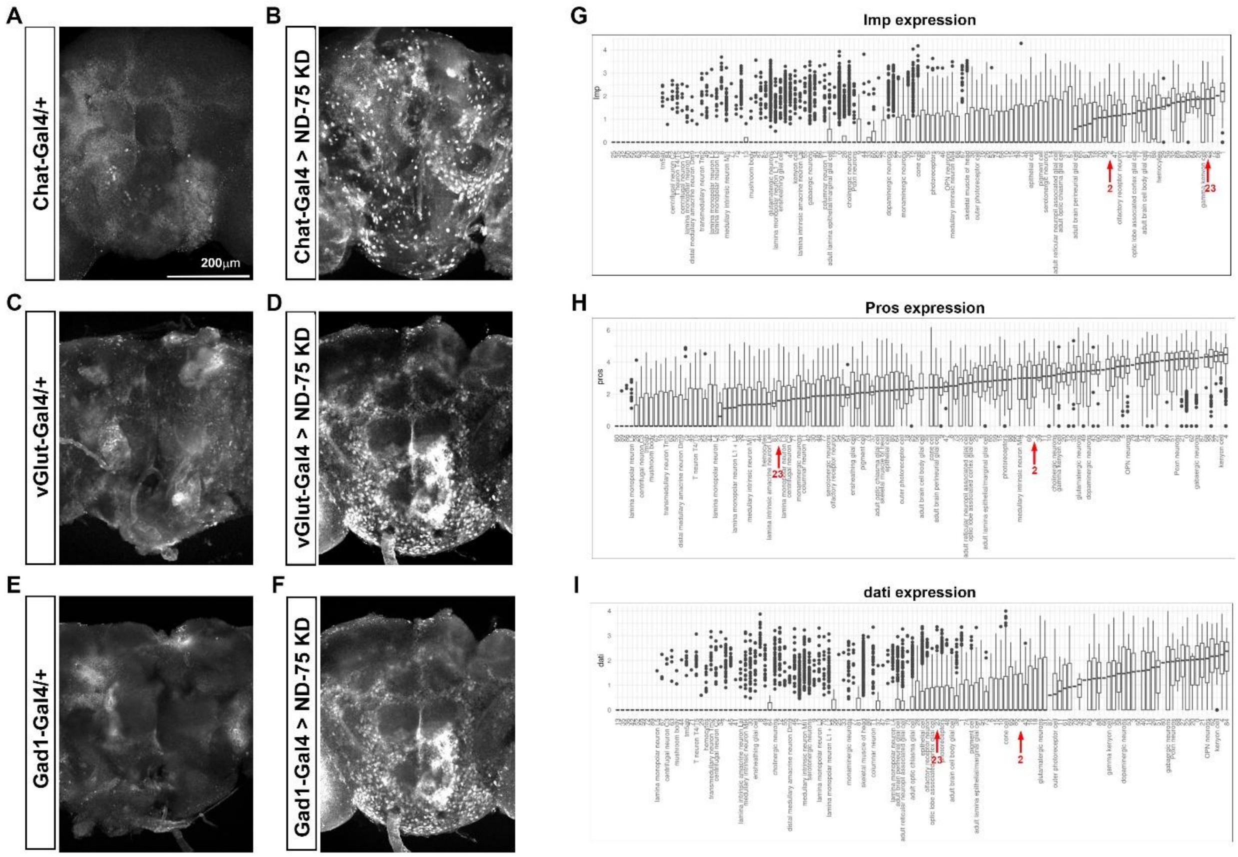


*Fig. S3. Novel clusters 2 and 23 have high expression of the central brain B neuron marker Imp but lower expression of the central brain A neuron markers Pros and dati*. (A-F). ND-75 KD in cholinergic neurons (using *Chat-Gal4*), glutamatergic neurons (using *vGlut-Gal4*) and GABAergic neurons (using *Gad1-Gal4*) activates ATF4. (G-I) Ranking of clusters by levels of *Imp* expression (G), *Pros* expression (E) and *dati* expression (F). Arrows indicate clusters 2 and 23.

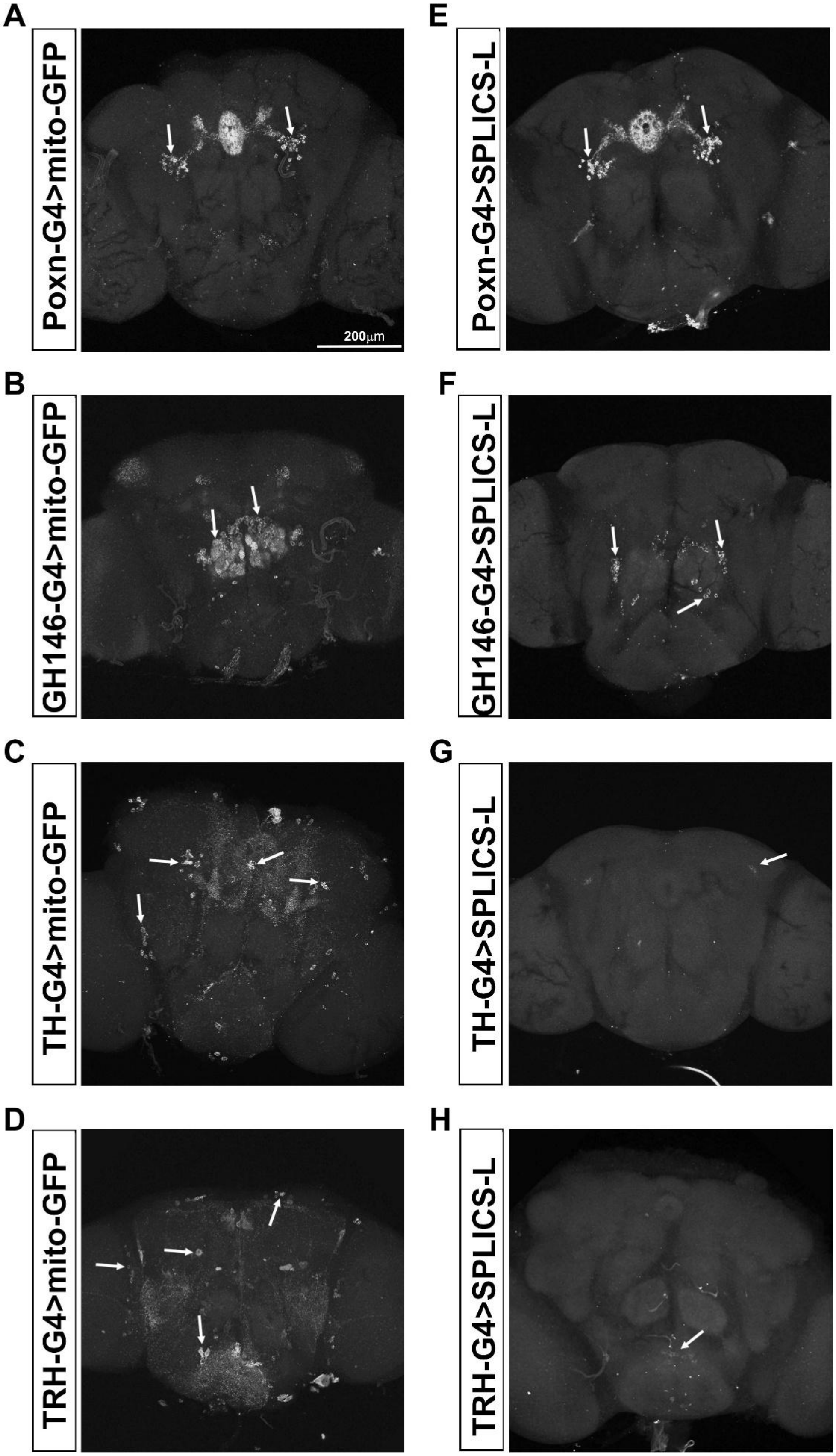
A
Poxn-G4>mito-GFP
200μm
B
GH146-G4>mito-GFP
C
TH-G4>mito-GFP
D
TRH-G4>mito-GFP
E
Poxn-G4>SPLICS-L
F
GH146-G4>SPLICS-L
G
TH-G4>SPLICS-L
H
TRH-G4>SPLICS-L

*Fig. S4. SPLICS-L, a reporter MERCs, is strongly expressed in Poxn neurons and OPNs but very weakly expressed in a few dopaminergic and serotonergic neurons.* (A-D) High levels of mito-GFP are visible in Poxn neurons (using *Poxn-Gal4*), OPNs (using *GH146-Gal4*), dopaminergic neurons (using *TH-Gal4*) and serotonergic neurons (using *TRH-Gal4*). Arrows indicate neuron cell bodies. (E, F) High levels of SPLICS-L are visible in cell bodies of Poxn neurons (using *Poxn-Gal4*) and OPNs (using *GH146-Gal4*). Arrows indicate neuron cell bodies. (G, H) Low levels of SPLICS-L are visible only in a few dopaminergic neuron cell bodies (using *TH-Gal4*) and serotonergic neuron cell bodies (using *TRH-Gal4*). Arrows indicate neuron cell bodies.

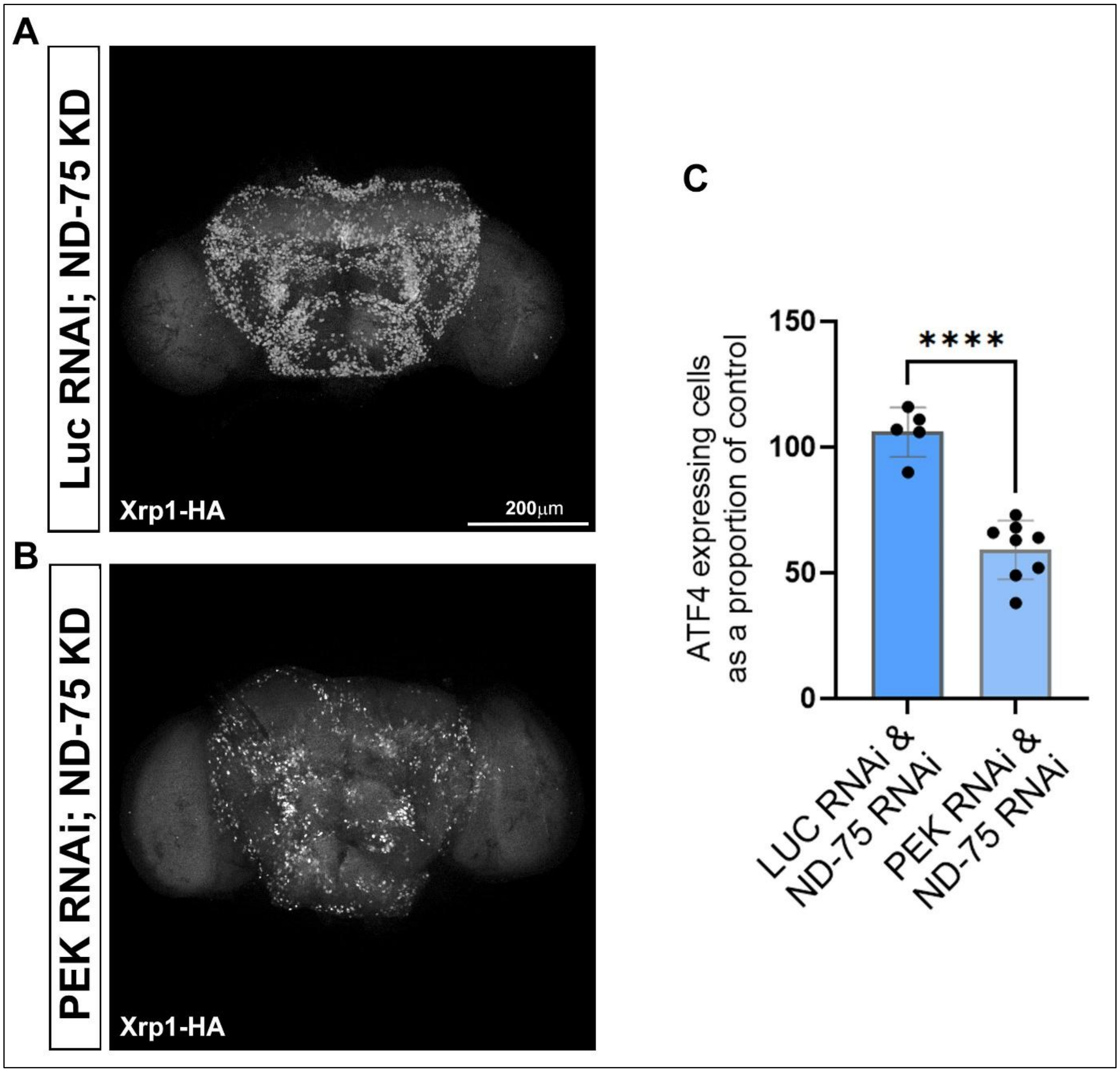


*Fig. S5. PEK is required for ND-75 KD-dependent activation of Xrp1.* (A, B) Pan-neuronal (*Tub-Gal80$^{ts}$;nSyb-Gal4*) ND-75 KD at 25ºC with luciferase RNAi (A) or PEK RNAi (B) shows that PEK knockdown reduces the number of neurons activating Xrp1-HA. (C) Quantification of Xrp1-HA activation by ND-75 KD with luciferase RNAi and PEK RNAi. Luc RNAi;ND-75 KD n=5 brains, PEK RNAi;ND-75 KD n=8 brains. Data are represented as mean ± SEM and were analysed using the student's unpaired t test. ****$p < 0.0001$.

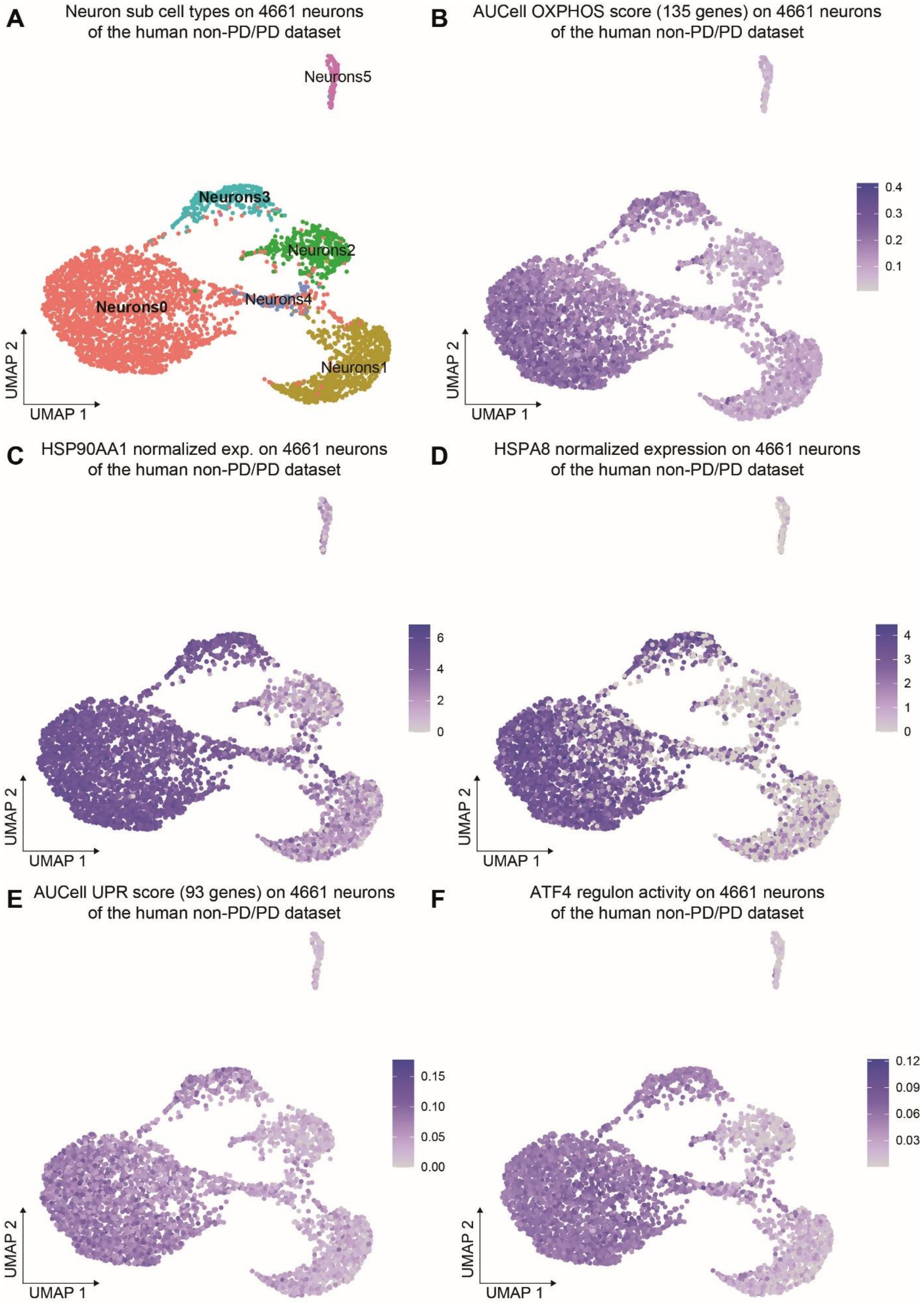
A
Neuron sub cell types on 4661 neurons of the human non-PD/PD dataset
Neurons5
Neurons3
Neurons2
Neurons4
Neurons0
Neurons1
UMAP 2
UMAP 1
B
AUCell OXPHOS score (135 genes) on 4661 neurons of the human non-PD/PD dataset
0.4
0.3
0.2
0.1
UMAP 2
UMAP 1
C
HSP90AA1 normalized exp. on 4661 neurons of the human non-PD/PD dataset
6
4
2
0
UMAP 2
UMAP 1
D
HSPA8 normalized expression on 4661 neurons of the human non-PD/PD dataset
4
3
2
1
0
UMAP 2
UMAP 1
E
AUCell UPR score (93 genes) on 4661 neurons of the human non-PD/PD dataset
0.15
0.10
0.05
0.00
UMAP 2
UMAP 1
F
ATF4 regulon activity on 4661 neurons of the human non-PD/PD dataset
0.12
0.09
0.06
0.03
UMAP 2
UMAP 1

*Fig. S6. OXPHOS gene expression correlates with UPR activation in the human SNpc*. (A) UMAP representation of 4,661 neurons extracted from the non-PD/PD dataset showing 6 clusters, labelled Neurons0-Neurons5. (B) UMAP representing AUCell aggregated score computed on 135 OXPHOS-related genes for 4,661 neurons extracted from the non-PD/PD dataset showing highest expression in Neurons0 and Neurons3 clusters. (C-F) UMAP representing AUCell aggregated score computed on *HSP90AA1* expression (C), *HSPA8* expression (D), 93 UPR-related genes (E) and SCENIC ATF4 regulon activity (F) in 4,661 neurons extracted from the non-PD/PD dataset showing highest OXPHOS, UPR and ATF4 activity in Neurons0 and Neurons3 clusters.

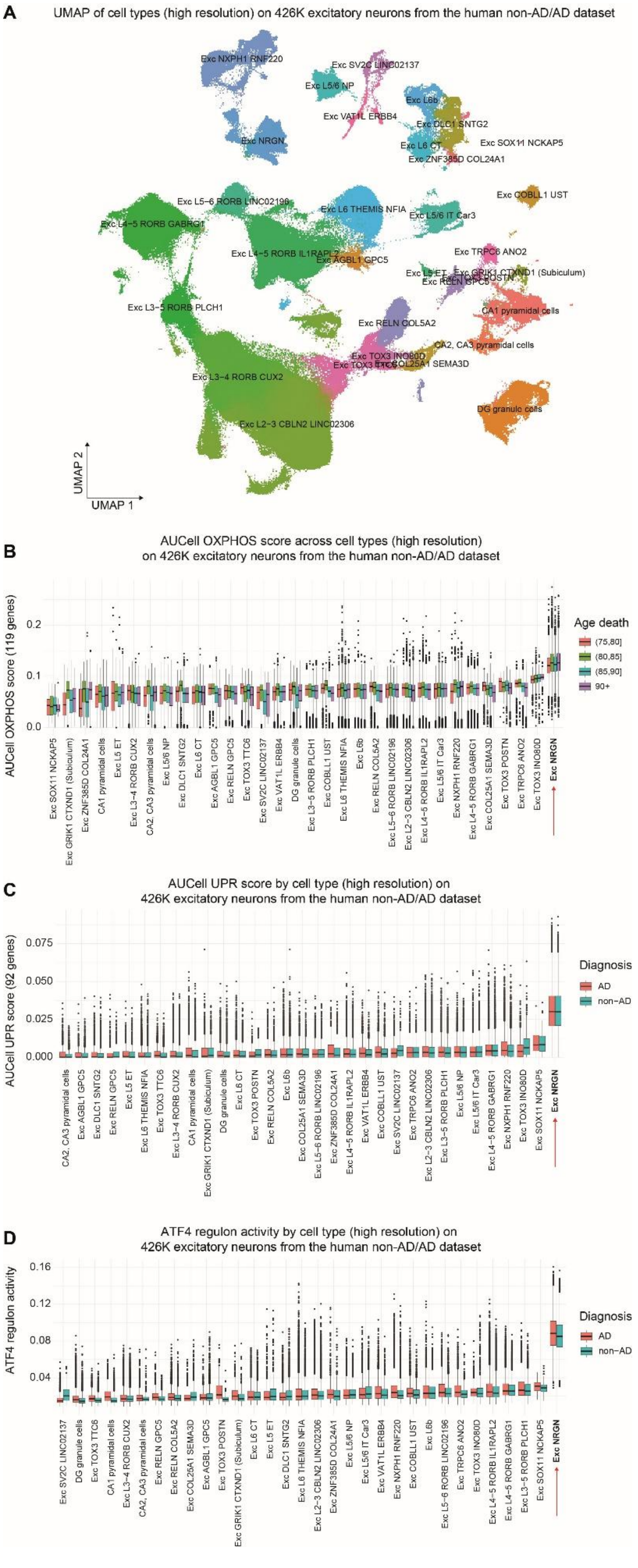
A
UMAP of cell types (high resolution) on 426K excitatory neurons from the human non-AD/AD dataset
Exc NXPH1 RNF220
Exc SV2C LINC02137
Exc L5/6 NP
Exc VAT1L ERBB4
Exc L6b
Exc DLC1 SNTG2
Exc L6 CT
Exc SOX11 NCKAP5
Exc ZNF385D COL24A1
Exc NRGN
Exc COBLL1 UST
Exc L5−6 RORB LINC02196
Exc L6 THEMIS NFIA
Exc L5/6 IT Car3
Exc L4−5 RORB GABRG1
Exc L4−5 RORB IL1RAPL2
Exc AGBL1 GPC5
Exc TRPC6 ANO2
Exc L3−5 RORB PLCH1
CA1 pyramidal cells
Exc RELN COL5A2
CA2, CA3 pyramidal cells
Exc L3−4 RORB CUX2
DG granule cells
Exc L2−3 CBLN2 LINC02306
UMAP 2
UMAP 1
B
AUCell OXPHOS score across cell types (high resolution)
on 426K excitatory neurons from the human non-AD/AD dataset
AUCell OXPHOS score (119 genes)
Age death
(75,80]
(80,85]
(85,90]
90+
Exc NRGN
C
AUCell UPR score by cell type (high resolution) on
426K excitatory neurons from the human non-AD/AD dataset
AUCell UPR score (92 genes)
Diagnosis
AD
non−AD
Exc NRGN
D
ATF4 regulon activity by cell type (high resolution) on
426K excitatory neurons from the human non-AD/AD dataset
ATF4 regulon activity
Diagnosis
AD
non−AD
Exc NRGN

*Fig. S7. Neurons with the highest OXPHOS gene expression activate the UPR in the human brain.* (A) UMAP representation of ~426k excitatory neurons extracted from the non-AD/AD dataset showing cell type annotations. (B-D) AUCell aggregated score computed on 119 OXPHOS-related genes (B), 92 UPR-related genes (C) and SCENIC ATF4 regulon activity (D) ranked for each excitatory neuron subtype of ~426k excitatory neurons extracted from the non-AD/AD dataset and split by patient death age range in (B), or patient clinical diagnosis (AD vs non-AD) in (C, D).

| Stock | Genotype | Source & stock number | Construct ID | Reference |
|---|---|---|---|---|
| $w^{1118}$ | w[1118] | BDSC 6326 | | |
| nSyb-Gal4 | y[1] w[*];P{nSyb-GAL4.S}3 | BDSC 51635 | | |
| Tubulin-$GAL80^{ts}$ | w[*]; P{w[+mC]=tubP-GAL80[ts]}10; TM2/TM6B, Tb[1] | BDSC 7108 | | |
| UAS-ND-75 RNAi | y[1] sc[*] v[1] sev[21];P{y[+t7.7] v[+t1.8]=TRiP.HMS00853}attP2 | BDSC 33910 | HMS00853 | [1] |
| UAS-PEK RNAi | y[1] v[1];UAS-PERK dsRNA | BDSC 42499 | HMJ02063 | |
| Luciferase RNAi | y[1] v[1];UAS-Luciferase dsRNA | BDSC 31603 | JF01355 | |
| Xrp1-HA | $Xrp1^{HA}$ | Nick Baker | | [2] |
| Imp-GFP | y[1]w[*] Mi{PT-GFSTF.2}Imp [MI05901-GFSTF.2] | BDSC 60237 | | |
| vGlut-Gal4 | P{w[+mC]=VGlut-GAL4.D}1, w[*] | BDSC 24635 | | |
| Gad1-Gal4 | P{w[+mC]=Gad1-GAL4.3.098}2/CyO | BDSC 51630 | | |
| Chat-Gal4 | w[*]; P{w[+mC]=ChAT-GAL4.7.4}19B/CyO | Alessio Vagnoni (BDSC 56500) | | |
| TH-Gal4 | y[1] w[1118]; TH-Gal4 | Frank Hirth | | [3] |
| TRH-Gal4 | y[1] w[1118]; TRH-Gal4/CyO | Frank Hirth | | [4] |
| Poxn-Gal4 | y[1] w[1118];Poxn-Gal4/CyO | Frank Hirth | | [5] |
| GH146-Gal4 | y[1] w[1118]; P{w[+mW.hs]=GawB}GH146 | BDSC 30026 | | |
| Unc84-GFP | w[*];UAS-Unc84-GFP (2) | Hongjie Li | | [6] |
| UAS-Opa1 | w[*]; P{w[+mC]=UAS-opa1.Flag3/TM6B, Tb[1] | BDSC 95258 | | |

| | | | | |
|---|---|---|---|---|
| *UAS-SPLICS-L* | *w[*];UAS-SPLICS*$_L$*/TM6B* | Alessio Vagnoni | | [7] |
| *UAS-SPLICS-S* | *w[*];UAS-SPLICS*$_s$*/CyO* | Alessio Vagnoni | | [8] |
| *UAS-mito-GFP* | *w[1118]; P{w[+mC]=UAS-mito-HA-GFP.AP}2/CyO* | BDSC 8442 | | [9] |
| *UAS-BiP-sfGFP-HDEL* | *w[1118]; PBac{y[+mDint2] w[+mC]=UAS-BiP-sfGFP-HDEL}VK00037* | BDSC 64748 | | |

*Table S1. Drosophila stocks genotype details.*

### Supplementary Data

*Data S1. Marker genes for cluster 2.*

*Data S2. Marker genes for cluster 23.*
